# Evaluating the effects of regularization and cross-validation parameters on the performance of SVM-based decoding of EEG data

**DOI:** 10.64898/2025.12.10.693548

**Authors:** Guanghui Zhang, Xinran Wang, Kurt Winsler, Steven J. Luck

**Author notes:** **Correspondence** Guanghui Zhang, Institute of Psychological and Brain Sciences, Liaoning Normal University, No. 850, Huanghe Road, Shahekou District, Dalian 116029, China. **Funding Information**. This study was made possible by grants R01MH087450 and R01EY033329 from the National Institutes of Health to Steven J. Luck; This study was also supported by Liaoning Normal University High-level Scientific Research Achievements Cultivation Project (No. 25GDL004), and the Natural Science Foundation of Liaoning Province (No. 2025-BS-0780) to Guanghui Zhang.

## Abstract

Regularization has been extensively used in multivariate pattern classification (MVPA; decoding) of EEG data to mitigate the risk of overfitting. N-fold cross-validation is also used to mitigate this risk, and it is often combined with averaging across trials to improve the SNR. However, the impact of different regularization and cross-validation parameters on decoding performance remains unclear. This study aimed to evaluate the effects of variations in the support vector machine (SVM) regularization parameter (C) and the number of crossfolds (and the number of trials per average) on the performance of SVM-based decoding analyses. To achieve this, we examined the decoding performance in relatively simple binary classification tasks from seven commonly used event-related potential paradigms (N170, mismatch negativity, N2pc, P3b, N400, lateralized readiness potential, and error-related negativity). Additionally, we evaluated the decoding performance in more challenging multiclass tasks, including decoding face identity, facial expression, stimulus location, and stimulus orientation. The results revealed that both decoding accuracy and effect size were highest when the regularization strength was equal to or greater than 1. Furthermore, using between 3 and 5 folds with at least 10 trials per average yielded optimal decoding performance in most cases. Researchers applying SVM-based decoding to datasets similar to those examined here—in terms of population, recording systems, class numbers, and paradigms—might benefit from using the parameters that we found to be optimal here.

## 1. Introduction

In recent decades, multivariate pattern classification (MVPA or *decoding*) has gained growing popularity in electroencephalogram (EEG) and event-related potential (ERP) research. This is due to the ability of MVPA to detect subtle differences between experimental conditions that traditional univariate approaches often fail to identify (Carrasco et al., 2024; Hebart & Baker, 2018; Peelen & Downing, 2023). For example, Carrasco et al. (2024) demonstrated that MVPA produced larger effect sizes across a wide range of ERP components compared to traditional univariate analyses. In addition, decoding analyses can handle noisier data more effectively because machine learning algorithms can focus on the most informative channels (Ashton et al., 2022; Carlson et al., 2019; Grootswagers et al., 2017). Here we will focus on classification via support vector machines (SVMs), which outperform other classification methods in many EEG datasets (Trammel et al., 2023), but we will also provide some analyses using linear discriminant analysis (LDA).

It is essential to deal with the problem of overfitting in MVPA so that the classification is based on consistent differences between classes rather than noise. In SVMs, one strategy for this is *regularization*, which helps control model complexity by penalizing large weights in the decision function. This reduces the risk of overfitting, especially when the number of features (e.g., EEG channels) is large relative to the number of trials (Hastie et al., 2009; Peng et al., 2021; Pereira et al., 2009). In many implementations of the SVM algorithm (e.g., Matlab’s fitcsvm function), the regularization strength is governed by the *box constraint parameter* (*C*). This parameter balances the trade-off between minimizing error in the training set and maintaining a simpler decision boundary that is more likely to generalize to new cases. A value of 1 gives equal weight to these two goals. A smaller *C* produces greater regularization, allowing more misclassifications in the training data in exchange for better generalization. By contrast, a larger *C* emphasizes correct classification of the training examples, potentially at the cost of overfitting. The use and reporting of regularization vary considerably across EEG/ERP decoding studies.

In the recent survey of the literature, a substantial proportion of studies did not explicitly mention whether regularization was applied or did not report the specific value of the regularization parameter (Xin et al., 2026). Many studies appeared to rely on the default settings implemented in common machine-learning toolboxes (e.g., the default *C* value in MATLAB’s *fitcsvm*() function) without further justification or tuning (e.g., Gomez-Andres et al., 2024; Sarrett & Toscano, 2024). By contrast, a smaller subset of studies explicitly applied and/or tuned regularization parameters (e.g., *C* = 0.1 versus 1; Den Ouden et al., 2025; Iamshchinina et al., 2022; Y. Li et al., 2025; Long et al., 2022). Additionally, a previous study showed that decoding accuracy across multiple ERP components (including N100, N200, P200, P300, and different LPP phases from one paradigm) improved substantially when the *C* parameter exceeded 0.1 (Tülay & Balli, 2024). This variability in reporting and implementation makes it difficult to evaluate the practical impact of regularization choices on decoding performance and highlights the need for a systematic examination of how regularization strength influences classification outcomes in multiple EEG/ERP datasets.

A second essential technique to address overfitting in EEG/ERP decoding is *N-fold cross-validation* (Lemm et al., 2011; Valente et al., 2021; Varoquaux, 2018; Varoquaux et al., 2017). This is often combined with averaging, which improves decoding performance by increasing the signal-to-noise ratio (SNR) of the data (Grootswagers et al., 2017; Petit et al., 2020; Williams et al., 2020, 2023). Specifically, after excluding trials with artifacts or other problems, the remaining trials are divided into *N* non-overlapping subsets, and the trials within each subset are averaged together to create pseudotrials. In every round of cross-validation, one pseudotrial from each class is used for testing, while the remaining pseudotrials are used for training. This process is repeated *N* times, with each pseudotrial serving as the test set once.

For example, one of the datasets analyzed by Carrasco et al. (2024) was an N170 experiment in which the two classes of interest were photographs of faces and photographs of cars, with 80 trials for each class (prior to the rejection of trials with errors or artifacts). A 5-fold leave-one-out cross-validation procedure was used, which created 5 averaged ERPs of approximately 16 trials for each class. In the first round of cross-validation, the first averaged ERP from each class was used as the test set, while ERPs 2–5 from each class served as the training set. In the second round, ERP 2 from each class was used as the test set and ERPs 1, 3, 4, and 5 were used as the training sets. This cycled three more times, using ERPs 3, 4, and 5 from each class as the test sets and the other ERPs as the training sets.

In this procedure, the number of crossfolds (*N*) is equivalent to the number of averaged ERPs per class, and the number of trials per ERP (*T*) is the number of total trials per class divided by *N* (and then rounded down to the nearest integer). The total number of trials per class is therefore *N*\**T*. Increasing the value of *N* will tend to increase decoding performance, all else being equal, because increasing *N* increases the number of training cases per class. However, increasing *N* reduces *T* and therefore decreases the SNR of each training and test case, and reducing *T* will tend to decrease decoding performance. A central goal of the present study was to determine what balance of *N* and *T* leads to the best decoding performance. Note that we did not consider any other cross-validation approaches (e.g., single-trial analyses), and we assume that the number of trials available for decoding is held constant such that *N* and *T* are reciprocally related to each other (as would be true in most real experiments, in which the total number of trials is fixed). We use the phrase *cross-validation parameters* to refer to the combination of *N* and *T*.

In current practice, *N* and *T* vary depending on the research topic and population, and they may even vary across studies within the same laboratory. The number of crossfolds (*N*) typically ranges from 2 to 25 or more, with the number of trials per average (*T*) typically varying between around 3 and 80 (Bae & Luck, 2018; Bayet et al., 2020; Bo et al., 2022; Chaisaen et al., 2020; Hong et al., 2020; Kikumoto & Mayr, 2018; Y. Li et al., 2022; Mares et al., 2020; Sanchez et al., 2020; Sarrett & Toscano, 2024; Turoman et al., 2024; Zhang, Carrasco, et al., 2024; Zhang, Wang, et al., 2025). A few previous studies have systematically varied the cross-validation parameters to assess their impact on decoding performance using real or simulated EEG datasets. For example, using real EEG data from a single study of auditory speech sounds, Sarrett et al. (2024) demonstrated that decoding accuracy improved as *T* increased and *N* decreased when they varied *T* from approximately 10 to 95 and *N* from 3 to 25 (Sarrett & Toscano, 2024).

Furthermore, a study using simulated EEG data suggested that decoding performance could be optimized by selecting *N* such that *T* is approximately 5-10% of the total number of available trials per class (Scrivener et al., 2023).

### 1.1 The goal of the current study

Because these previous studies of the cross-validation parameters each focused on a single dataset (Sarrett & Toscano, 2024; Scrivener et al., 2023), it is difficult to know whether their conclusions would generalize to datasets employing different experimental paradigms, electrode densities, and numbers of stimulus classes. The goal of the present study was to provide a more systematic and generalizable assessment of the impact of the cross-validation parameters and the regularization strength on SVM-based EEG/ERP decoding performance. We accomplished this by using a wide range of EEG/ERP experiments with diverse sample sizes, a wide range of numbers of stimulus classes, and different electrode densities.

We analyzed the data from multiple publicly available EEG/ERP datasets. One of these datasets was the ERP CORE (Compendium of Open Resources and Experiments; Kappenman et al., 2021), which includes six common ERP paradigms that can be used to elicit seven distinct ERP components: N170, mismatch negativity (MMN), P3b, N400, error-related negativity (ERN), N2pc, and lateralized readiness potential (LRP). For these experiments, binary decoding was applied to the primary classes (e.g., faces versus cars for the N170 experiment, standard versus deviant auditory stimuli for the MMN experiment). These datasets consist of recordings from 32 electrode sites with a moderate density across the scalp.

We also analyzed three additional datasets that involved more challenging multiclass decoding tasks, where the differences among classes were difficult to detect using conventional univariate analyses. In one of the paradigms, the stimulus set consisted of 16 different photographs of faces, combining 4 face identities with 4 emotional expressions (Bae, 2021). We decoded face identity while disregarding facial expression and facial expression while disregarding face identity. For this dataset, recordings were obtained from 64 electrodes, providing greater electrode density than in the other datasets.

In the other two paradigms, participants were required to perceive and remember one of 16 different orientations, and data were recorded from 32 electrodes (Bae & Luck, 2018). In one of these experiments, there were also 16 different stimulus locations. For these two paradigms, we decoded the data in the frequency domain as well as in the time domain (as in the original report of these datasets).

When evaluating the impact of the regularization and cross-validation parameters on decoding, it is important to ask what should be considered “better” decoding performance. In the present analyses, we are focusing on datasets that address scientific goals (e.g., determining how the brain works, understanding the nature of brain disorders) rather than engineering goals (e.g., building brain-computer interfaces). As discussed by Hebart and Baker (2018), these goals have different requirements. Whereas real-time single-trial accuracy is important for most engineering goals, scientific goals can often be achieved by averaging across trials and identifying small effects, as long as these effects are statistically significant and theoretically meaningful. The present study therefore seeks to find the decoding parameters that yield the greatest sensitivity for distinguishing among the experimental conditions being classified. Greater decoding accuracy will often be an index of greater sensitivity, and we therefore ask which parameters yield the greatest decoding accuracy. However, if greater decoding accuracy comes at the cost of greater variability in decoding accuracy across participants, then it will not necessarily yield greater statistical power for determining whether decoding is above chance or differs across groups or experimental conditions. We therefore also quantify the *effect size* of the decoding accuracy, which scales the decoding accuracy by the variability across participants. Effect size is directly related to statistical power, so parameters that maximize the effect size in a study will also generally maximize the statistical power of the study.

## 2. Methods

The ERP CORE dataset was downloaded from https://osf.io/thsqg/, and the other datasets were downloaded from https://osf.io/tgzew/ or https://pan.baidu.com/s/1VCZ7YERDRfqAFvjIgmYGFA?pwd=1234. The data analysis procedures were implemented with MATLAB 2021b (MathWorks Inc), using EEGLAB Toolbox v2023.0 (Delorme & Makeig, 2004) combined with ERPLAB Toolbox v11.00 (Lopez-Calderon & Luck, 2014). The processing scripts are available at https://osf.io/un6pq.

The following sections provide a brief overview of the stimuli, tasks, recording methods, and preprocessing techniques for each dataset. Comprehensive descriptions are available in the original papers that detail these datasets, which we referred to as the *ERP CORE* (Kappenman et al., 2021), *Orientations* (Bae & Luck, 2018), and *Faces* (Bae, 2021) datasets. All the studies were approved by the UC Davis Institutional Review Board.

### 2.1 Participants

All the participants were highly cooperative neurotypical college students from the UC-Davis community. The number of participants with usable data for the ERP CORE, Orientations, and Faces experiments is shown in Table 1.

**Table 1.** Epoch window, baseline period, electrode site, number of classes, and measurement window used for each experiment. Note that we collapsed the 16 stimuli into 4 classes for the Orientations-1 and Orientation-2 experiments in our analyses of the effects of the cross-validation parameters.

| Experiment | Epoch (ms) | Baseline Period (ms) | Measurement window (ms) | # of Subjects | # of classes (M) | # of channels |
| --- | --- | --- | --- | --- | --- | --- |
| N170 | -200 to 800 | -200 to 0 | 110 to 150 | 39 | 2<br>(faces vs. cars) | 28 |
| MMN | -200 to 800 | -200 to 0 | 125 to 225 | 40 | 2<br>(deviants vs. standards) | 28 |
| P3b | -200 to 800 | -200 to 0 | 300 to 600 | 38 | 2<br>(targets vs. nontargets) | 28 |
| N400 | -200 to 800 | -200 to 0 | 300 to 500 | 39 | 2 (related vs. unrelated) | 28 |
| ERN | -600 to 400 | -400 to -200 | 0 to 100 | 22 | 2<br>(correct vs. error) | 28 |
| LRP | -800 to 200 | -800 to -600 | -100 to 0 | 40 | 2<br>(left hand vs. right hand) | 28 |
| N2pc | -200 to 800 | -200 to 0 | 200 to 275 | 40 | 2<br>(left target vs. right target) | 28 |
| Orientations-1 | -500 to 1500 | -500 to 0 | 100 to 500<br>or<br>500 to 1000 | 16 | 4 or 16<br>(orientation classes) | 27 |
| Orientations-2 | -500 to 1500 | -500 to 0 | 100 to 500<br>or<br>500 to 1000 | 16 | 4 or 16<br>(orientation or location classes) | 27 |
| Faces | -500 to 1500 | -500 to 0 | 100 to 500<br>or<br>500 to 1000 | 22 | 4 (face identities or emotional expressions) | 59 |

**Table 2.** Number of trials for each condition being decoding for the ERP CORE, Orientations-1 & -2, and Faces datasets. Note that, for the “Orientation” studies, each row corresponds to a distinct orientation/location condition rather than a different study.

| ERP CORE |  |  | Orientation 1 |  | Orientation 2 |  |  |  | Face |  |
| --- | --- | --- | --- | --- | --- | --- | --- | --- | --- | --- |
| ERP component | condition | Mean ± SD | 16 stimuli | 4 stimuli | Location | | Orientation | | Facial ID | Facial expression |
|  |  |  |  |  | 16 stimuli | 4 stimuli | 16 stimuli | 4 stimuli |  |  |
| P3 | Rare | 34.92 ± 4.49 | 39.00 ± 0.65 | 118.47 ± 0.64 | 40.00 ± 0.00 | 120.00 ± 0.00 | 40.00 ± 0.00 | 120.00 ± 0.00 | 161.36 ± 6.40 | 160.82 ± 3.84 |
| | Frequent | 154.39 ± 6.61 | 39.53 ± 0.52 | | 40.00 ± 0.00 | | 40.00 ± 0.00 | | | |
| N170 | Faces | 75.13 ± 5.27 | 39.33 ± 0.82 | | 40.00 ± 0.00 | | 40.00 ± 0.00 | | | |
| | Cars | 72.28 ± 6.61 | 39.60 ± 0.51 | | 40.00 ± 0.00 | | 40.00 ± 0.00 | | | |
| MMN | Deviants | 199.18 ± 5.22 | 39.13 ± 0.74 | 118.67 ± 0.62 | 40.00 ± 0.00 | 120.00 ± 0.00 | 40.00 ± 0.00 | 120.00 ± 0.00 | 160.50 ± 2.58 | 160.95 ± 4.71 |
| | Standards | 581.75 ± 15.56 | 39.53 ± 0.74 | | 40.00 ± 0.00 | | 40.00 ± 0.00 | | | |
| N400 | Unrelated | 56.62 ± 3.34 | 39.47 ± 0.52 | | 40.00 ± 0.00 | | 40.00 ± 0.00 | | | |
| | related | 56.38 ± 2.62 | 39.67 ± 0.49 | | 40.00 ± 0.00 | | 40.00 ± 0.00 | | | |
| ERN | Incorrect | 59.23 ± 13.01 | 39.00 ± 1.00 | 117.73 ± 1.10 | 40.00 ± 0.00 | 120.00 ± 0.00 | 40.00 ± 0.00 | 120.00 ± 0.00 | 161.09 ± 5.12 | 161.32 ± 6.18 |
| | Correct | 330.36 ± 36.13 | 39.40 ± 0.83 | | 40.00 ± 0.00 | | 40.00 ± 0.00 | | | |
| N2pc | Left | 138.65 ± 11.54 | 38.87 ± 0.52 | | 40.00 ± 0.00 | | 40.00 ± 0.00 | | | |
| | Right | 141.10 ± 12.91 | 39.47 ± 0.52 | | 40.00 ± 0.00 | | 40.00 ± 0.00 | | | |
| LRP | Left | 175.53 ± 19.65 | 39.27 ± 0.46 | 118.67 ± 0.6172 | 40.00 ± 0.00 | 120.00 ± 0.00 | 40.00 ± 0.00 | 120.00 ± 0.00 | 161.09 ± 5.12 | 160.95 ± 4.47 |
| | | | 39.27 ± 0.59 | | 40.00 ± 0.00 | | 40.00 ± 0.00 | | | |
| | Right | 176.55 ± 17.89 | 39.53 ± 0.64 | | 40.00 ± 0.00 | | 40.00 ± 0.00 | | | |
| | | | 39.87 ± 0.35 | | 40.00 ± 0.00 | | 40.00 ± 0.00 | | | |

### 2.2 Paradigms

Each of the ERP CORE paradigms lasted approximately 10 minutes, with each participant completing all six tasks in a single session (Kappenman et al., 2021). For the N170 face perception task (Figure 1a), participants were instructed to categorize each stimulus as a "texture" (scrambled faces or scrambled cars) or an "object" (faces or cars). Here we analyzed data only from face and car stimuli. As shown in Figure 1b, a passive auditory oddball task was used to elicit the mismatch negativity (MMN). During the task, participants heard a task-irrelevant sequence of standard (80 dB) and deviant (70 dB) auditory tones while viewing a silent video. The P3b component (Figure 1c) was elicited using an active visual oddball task, in which participants made one response to target letters and a different response to non-target letters. In each block, one of five letters (A, B, C, D, E; p = .2 each) was designated the target and the remaining letters served as non-targets. The N400 component (Figure 1d) was elicited using a word-pair judgment task. Each trial consisted of a prime word and a target word, and participants were required to determine whether the target word was semantically related to the prime word (p = .5 for related and unrelated). The lateralized readiness potential (LRP) and error-related negativity (ERN; Figure 1e) were elicited via an Eriksen flankers task, in which participants responded to the direction of a central arrowhead surrounded by congruent or incongruent arrowheads by pressing corresponding buttons with their left or right hand. The N2pc component (Figure 1f) was elicited during a visual search task, in which 24 colored squares were presented on each trial and participants reported the location (top or bottom) of a gap in the one square drawn in a designated target color (pink or blue). In all tasks, participants were instructed to maintain fixation on a central point. In all but the MMN paradigm, participants pressed one of two buttons on each trial to report the category of the current trial stimulus.

**Figure 1.**
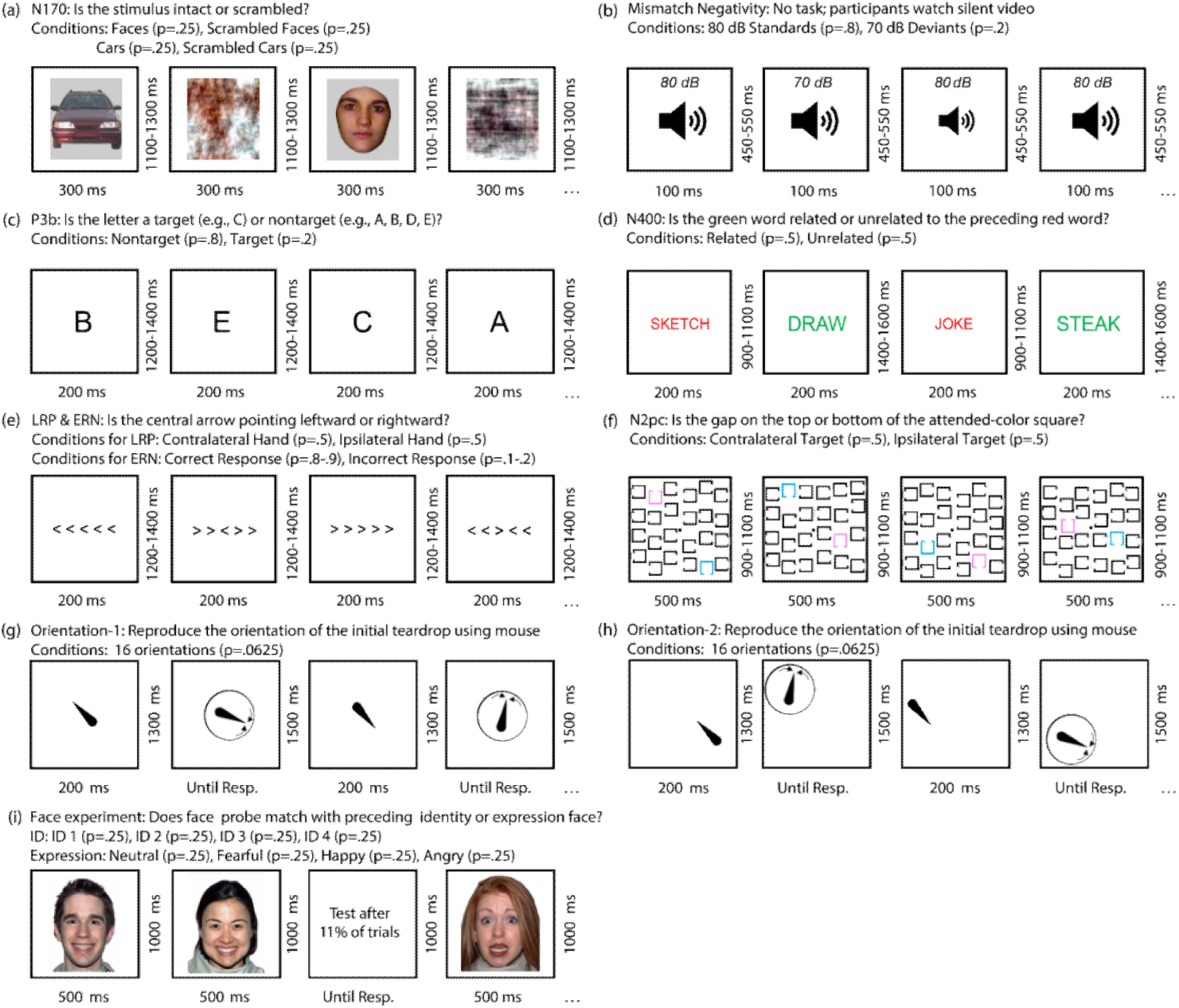
Example trials from the experimental paradigms. (a) N170 Paradigm: Participants identified whether a given stimulus was intact (faces or cars) or scrambled (scrambled faces or cars). This study focused exclusively on the face and car trials. (b) Mismatch Negativity Paradigm: Participants watched a silent video while task-irrelevant standard tones (80 dB, probability = .8) and deviant tones (70 dB, probability = .2) were presented. (c) P3b Paradigm: The letters A, B, C, D, and E were presented in random order, each with a probability of .2. One letter was designated as the target for each trial block, and participants identified whether a given stimulus was the target or a nontarget. (d) N400 Paradigm: Participants determined whether a green target word was semantically related or unrelated to a preceding red prime word. (e) Flankers Task: This task elicited the lateralized readiness potential (LRP) and error-related negativity (ERN). Participants reported the direction of the central arrow (leftward or rightward) while ignoring the surrounding arrows. (f) N2pc Paradigm: For each trial block, either pink or blue was designated as the target color. Participants reported whether the gap on the target-colored square was at the top or bottom of the square. (g) Orientations-1 Paradigm: Participants memorized the orientation of an initial teardrop, retained it over a 1300-ms delay period, and then used a mouse to adjust the orientation of a test teardrop to match the original. (h) Orientations-2 Paradigm: Identical to Orientations-1 except the teardrop appeared randomly at one of 16 different locations. Note that the orientations or locations located in the same quadrant are combined to one new orientation or location. (i) Face Paradigm: In each trial, one of 16 faces was presented, representing a combination of four unique identities and four distinct emotional expressions. On 11% of the trials, participants were asked to recall the most recent face shown.

In the Orientations-1 paradigm (Bae & Luck, 2018; Figure 1g), a teardrop shape (the *sample* stimulus) appeared at the center of the display for 200 ms at the start of each trial. The teardrop could take one of 16 possible orientations, spaced 22.5° apart. After a 1300-ms blank delay, participants adjusted the orientation of a *test* teardrop using a mouse to match the remembered orientation of the sample teardrop. In the Orientations-2 paradigm (Figure 1h), the sample teardrop was again presented in one of 16 random orientations, but its location varied across 16 possible positions around an invisible circle. The orientation and location were independent of each other. After the delay period, participants reproduced the teardrop’s orientation with a test teardrop in a random location, making the location task-irrelevant.

The Faces dataset comes from a single experiment (Figure 1i) (Bae, 2021). On each trial, a sample face was first presented at the center of the display for 500 ms, followed by a 1000-ms blank interval. The face was randomly chosen from a set of 16 combinations of four identities (ID 1, ID 2, ID 3, and ID 4) and four emotional expressions (neutral, fearful, happy, and angry). On 89% of trials, the delay was followed by the next face stimulus. On the remaining 11% of trials, a probe display with four faces appeared. Participants were required to select the face in the probe that matched the immediately preceding sample face in either identity or expression. Identity and expression probes were presented randomly, requiring participants to remember both aspects of each face.

### 2.3 EEG recording and preprocessing

For the ERP CORE paradigms, the EEG was recorded using a Biosemi ActiveTwo system (Biosemi B.V.) at a sampling rate of 1024 Hz. During acquisition, a fifth-order sinc filter with a half-power cutoff at 204.8 Hz was applied to prevent aliasing. The dataset contains EEG signals from 30 scalp electrodes: FP1/2, Fz, F3/4/7/8, FCz, FC3/4, Cz, C3/4/5/6, CPz, Pz, P3/4/7/8, PO3/4/7/8/9/10, and Oz/1/2. The dataset also contains signals from two electrooculogram (EOG) electrodes placed lateral to the eyes and one EOG electrode below the right eye.

For the Faces and Orientations paradigms, the EEG was recorded using a Brain Products actiCHamp system (Brain Products GmbH) with a sampling rate of 500 Hz. To avoid aliasing, a cascaded integrator-comb filter was applied during recording, with a half-power cutoff at 130 Hz. These datasets included the same EOG signals as in the ERP CORE dataset, as well as EEG electrodes placed at the left and right mastoids. For both Orientations paradigms, EEG was collected from 27 EEG sites: FP1/2, Fz, F3/4/7/8, Cz, C3/4, Pz, P3/4/5/6/7/8/9/10, POz, PO3/4/7/8, and Oz/1/2. The Faces dataset includes 59 EEG sites: FP1/2, AFz/3/4/7/8, Fz/1/2/3/4/5/6/7/8, FCz/1/2/3/4/5/6, Cz/1/2/3/4/5/6, T7/8, CPz/1/2/3/4/5/6, TP7/8, Pz/1/2/3/4/5/6/7/9/10, POz/3/4/7/8, Oz/1/2.

The key preprocessing steps for the ERP CORE paradigms are provided below, and more detailed descriptions can be found in the original paper (Kappenman et al., 2021). The continuous EEG signal was downsampled to 256 Hz and referenced to the average of the P9 and P10 electrodes (located near the left and right mastoids). However, in the N170 paradigm, the average of all scalp sites served as the reference. A noncausal Butterworth bandpass filter (half amplitude cutoff = 0.1 – 30 Hz, roll-off = 12 dB/oct) was applied. Note that subsequent research has demonstrated that different filter settings should be used for each paradigm to optimize statistical power for univariate analyses (Zhang, Garrett, & Luck, 2024b, 2024a), but we held the filter settings constant across paradigms here to avoid introducing a confound in comparing the different paradigms. To correct for blinks, independent component analysis (ICA; T. Jung et al., 2000; T.-P. Jung et al., 2000) was employed. Instead of the original ICA procedure of ERP CORE paradigms, we used an optimized ICA procedure that has been shown to be more effective (Dimigen, 2020; Klug & Gramann, 2021; Luck, 2022; Zhang, Garrett, Simmons, et al., 2024). In this procedure, the ICA decomposition is performed on a copy of the dataset that has been heavily filtered (1-30 Hz) and then the weights are transformed back to the original dataset prior to performing the correction. After the correction, the datasets were segmented into epochs and baseline-corrected using the time windows shown in Table 1. Channels identified as problematic were interpolated using ERPLAB’s spherical spline interpolation method. The decoding analysis included all trials except those with behavioral errors, because we have shown that the rejection of trials containing artifacts has little impact on decoding performance across a broad range of EEG and ERP paradigms (Zhang & Luck, 2025).

The remaining preprocessing steps for the Faces and Orientations paradigms mirrored those of the ERP CORE, with the exception that the data were downsampled to 250 Hz and referenced to the average of the mastoids. In addition, no trials were excluded because of behavioral errors, which were very rare. For further details, refer to Bae (2021) and Bae and Luck (2018), and consult Table 1 for epoch timing.

### 2.4 SVM-based decoding methods

We performed decoding multiple times for each dataset, each time using one of several different regularization or cross-validation parameters. We performed binary decoding for the ERP CORE paradigms. In the N170, MMN, P3b, and N400 paradigms, we decoded which of two stimulus classes was presented (e.g., targets versus nontargets in the P3b paradigm). In the N2pc paradigm, we decoded whether the target was on the left or right side of the display. In the flankers paradigm, we decoded whether the participant responded with the left hand or the right hand for the LRP analysis (for correct responses), and we decoded whether the response was correct or incorrect for the ERN analysis (disregarding whether the left or right hand was used to respond).

For the Faces dataset, we performed two independent 4-class decoding runs for each set of regularization and cross-validation parameters. In the first run, we decoded which of the four face identities was present, disregarding the emotional expression of the face. In the second run, we decoded which of the four emotional expressions was displayed, disregarding the identity of the face.

When we manipulated the regularization parameter for the Orientations experiments, we classified which of the 16 orientations was presented, disregarding stimulus location. For Orientations-2, we also decoded which of the 16 stimulus locations was used, disregarding orientation. We modified the approach somewhat to test the cross-validation parameters in the Orientations experiments. There were only 40 trials per orientation or per location, which made it difficult to examine a broad set of *T* and *N* values. To assess the effects of the cross-validation parameters, we therefore collapsed the 16 orientations or the 16 locations into 4 classes (upper left, upper right, lower left, and lower right) and decoded which of these 4 orientation classes or location classes was present. This yielded 120 trials per class.

We decoded the data separately at each time point for each participant in each experiment, using the set of EEG channels as the feature set for decoding. For binary decoding, we used the MATLAB function *fitcsvm()* to train the linear SVM classifier. For multiclass decoding, we used the MATLAB function *fitcecoc()* to implement the error-correcting output codes approach. After training, we used the MATLAB function *predict()* to compute decoding accuracy for the test data.

In our leave-one-out cross-validation approach, we first split the available trials for a given participant into *N* subsets or folds for each stimulus class, resulting in *T* trials per subset where T is equal to the total number of available trials per class divided by *N* (Hou et al., 2025, 2026; Song et al., 2025; Zhang, Xin, et al., 2025). We then averaged the T trials for each subset to create *N* averaged ERPs for each class. For the data at a specific time point, we used *N-1* of the averaged ERPs from each class to train the SVM decoder, and we then tested the decoder with the remaining averaged ERP from each class.

We repeated this procedure *N* times, with each set of averaged ERPs used as the test set one time and as a training set N-1 times. The total number of tests was therefore *N*K* for each repetition of the procedure, where *K* is the number of classes being decoded. We then iterated this whole process J times, resulting in a proportion correct over a total of *J*K*N* tests. The key factor in controlling the precision of a proportion value is the total number of tests. For each paradigm, we chose a number of iterations (*J*) that yielded 5000 tests. This held the precision of the proportion correct variable constant even though the number of classes (*K*) varied across paradigms.

When the total available trials varied across classes (e.g., in experiments using oddball paradigms), we subsampled from the classes as necessary to equate *T* across classes. This subsampling is essential, because the SNR must be equal across classes to avoid spurious above-chance decoding. That is, if the pseudotrials are noisier for one class than for the other class(es), the pseudotrials for the noisier class are more likely to contain extreme values, and the decoder can perform better than chance by selecting a relatively extreme decision hyperplane. Equating *T* across classes is therefore necessary.

To examine the effect of the regularization strength (the box constraint parameter C), we performed the decoding procedure for a given dataset once for each of several values of C (0.001, 0.01, 0.1, 1, 10, 100, and 1000). For these analyses, we used an N of 3 for the N-fold cross-validation procedure, with the maximum number of trials per average (T) that was possible for that N in a given dataset (see the supplementary materials for evidence that variations in C produce similar effects whether N is 20 or 3). By holding N constant, we could ensure that any differences in decoding performance across regularization parameters reflected the effect of regularization rather than the effects of the cross-validation parameters.

To test the effects of the N and T cross-validation parameters, we held the regularization strength constant at a balanced level (C = 1), and we repeated the decoding procedure for each dataset once with each of several values of N and T. Specifically, we examined N = 2, 3, 4, 5, 6, 10, 20, and 40 crossfolds for each paradigm (except that we could not test 40 crossfolds for the P3b paradigm because the total number of trials for one condition was smaller than 40), with T set to the highest value possible given that value of N for a given dataset.

Bae and Luck (2018) found that location and orientation in the Orientations datasets could be decoded from the alpha-band EEG power as well as from the ERP amplitude. We therefore decoded each of these datasets using both alpha-band power and ERP amplitude (separately). The alpha-band decoding was identical to the ERP decoding, except that we applied a bandpass filter from 8-12 Hz (two-way least-squares finite impulse response filter) and passed the filtered data through the Hilbert transform to extract the phase-independent power in this frequency range.

### 2.5 Time-window averaging and effect size calculation

For the sake of simplicity, decoding accuracy was averaged within a time window that was defined a priori for each paradigm (see Table 1). For the ERP CORE experiments, we used the same time windows that were used for the original univariate analyses of each experiment. For the other paradigms, we used both a perceptual time window (100–500 ms) and a working memory time window (500–1000 ms), as in previous methodology studies using these datasets (Bae & Luck, 2018).

Although researchers often try to maximize decoding accuracy, it is also important to consider statistical power, which is influenced by both the mean decoding accuracy across participants and variability across participants. We therefore also examined the Cohen’s dz metric of effect size, which is directly related to statistical power and reflects both mean decoding accuracy and participant variability. We calculated Cohen’s dz for each analysis as the mean across participants divided by the standard deviation across participants (after first subtracting the chance level for a given analysis from each participant’s decoding accuracy). Bootstrapping with 10,000 iterations was employed to estimate the standard error of this metric of effect size.

In many cases, researchers look at decoding accuracy separately at each time point rather than averaging across a time window. Taking the mean decoding accuracy across a time window of interest captures the central tendency of the single-point decoding accuracy values during that window. However, the effect size will be larger when computed from the average decoding accuracy across a time window than when computed individually at each time point. To characterize the central tendency of the single-point effect sizes, we also computed the effect size separately at each time point and then averaged those effect size values over the time range of interest (see supplementary Figure S11).

### 2.6 Statistical analyses

To determine whether the effects of the regularization and cross-validation parameters had a statistically significant impact on decoding accuracy, we conducted mixed-effects regression analyses for each set of decoding accuracy results. For each data set, we used the R package lme4 (Bates et al., 2015) to fit two linear mixed-effect regression models; one predicting decoding accuracy as a function of the regularization parameter C (log₁₀-transformed), and another as a function of the number of trials (log₁₀-transformed) used per fold. Because some of the effects were clearly nonmonotonic, both linear and quadratic terms for the predictor were included as fixed effects. Predictors were centered and scaled (*z*-scored) prior to fitting. Random intercepts were included for subjects, and we attempted to fit correlated random slopes for each of the predictor terms. If a model had a singular fit or failed to converge, we simplified the random effect structure first by removing the correlation between the quadratic slope and intercept, followed by removing the random slope for the quadratic term if needed. Most models were able to be estimated with the full random effect structure. Significance of the linear and quadratic terms was assessed via *t*-tests with Satterthwaite-approximated degrees of freedom using the lmerTest package (Kuznetsova et al., 2017). The joint significance of both predictors together was tested via a likelihood ratio test comparing the full model to an intercept-only fixed-effects model with the same random effects structure. The results are shown in Table 3.

**Table 3.** Linear and quadratic regression results.

| Experiment |  | Measurement windows(ms) | Regularization |  |  |  | Crossfold |  |  |  |
| --- | --- | --- | --- | --- | --- | --- | --- | --- | --- | --- |
|  |  |  | Linear effect |  | Quadratic effect |  | Linear effect |  | Quadratic effect |  |
|  |  |  | t | p | t | p | t | p | t | p |
| N170 |  | 110 to 150 | 2.860 | 0.007 | 0.116 | 0.908 | 8.712 | **<0.001** | -5.922 | **<0.001** |
| MMN |  | 125 to 225 | 8.178 | **<0.001** | -7.664 | **<0.001** | 7.641 | **<0.001** | 0.201 | 0.842 |
| P3b |  | 300 to 600 | 0.121 | 0.904 | -0.739 | 0.464 | 9.856 | **<0.001** | -6.960 | **<0.001** |
| N400 |  | 300 to 500 | -0.495 | 0.623 | -3.101 | 0.003 | 14.568 | **<0.001** | -6.049 | **<0.001** |
| ERN |  | 0 to 100 | 5.770 | **<0.001** | -5.575 | **<0.001** | 7.746 | **<0.001** | -4.847 | **<0.001** |
| LRP |  | -100 to 0 | 1.270 | 0.212 | -1.480 | 0.147 | 1.813 | 0.078 | -12.366 | **<0.001** |
| N2pc |  | 200 to 275 | 1.041 | 0.304 | -1.879 | 0.068 | 1.672 | 0.103 | -6.432 | **<0.001** |
| Orientations-1 | 0.1-30Hz | 100 to 500 | 1.960 | 0.070 | -3.039 | 0.003 | 0.304 | -7.966 | **<0.001** | **<0.001** |
|  |  | 500 to 1000 | -1.255 | 0.230 | 0.672 | 0.512 | 0.912 | -2.887 | **0.011** | **0.002** |
|  | Alpha | 100 to 500 | 0.381 | 0.709 | -0.771 | 0.454 | 0.471 | -3.516 | **0.003** | **<0.001** |
|  |  | 500 to 1000 | -1.049 | 0.312 | 1.779 | 0.097 | **0.003** | -3.223 | **0.006** | **<0.001** |
| Orientations-2 | 0.1-30Hz | 100 to 500 | 0.105 | 0.918 | -1.148 | 0.269 | 0.451 | -4.450 | **<0.001** | **<0.001** |
|  |  | 500 to 1000 | 1.100 | 0.289 | -2.607 | 0.020 | 0.215 | -1.858 | 0.083 | **0.001** |
|  | Alpha | 100 to 500 | -0.704 | 0.492 | -0.619 | 0.545 | 0.362 | -0.129 | 0.899 | 0.322 |
|  |  | 500 to 1000 | -1.988 | 0.065 | 1.492 | 0.156 | 0.392 | -0.419 | 0.681 | 0.193 |
| Location | 0.1-30Hz | 100 to 500 | -0.299 | 0.769 | -1.309 | 0.210 | **0.006** | -6.654 | **<0.001** | **<0.001** |
|  |  | 500 to 1000 | -1.263 | 0.226 | -0.458 | 0.653 | **0.036** | -3.304 | **0.005** | **<0.001** |
|  | Alpha | 100 to 500 | -0.679 | 0.508 | -0.105 | 0.918 | **0.001** | -3.114 | **0.007** | **<0.001** |
|  |  | 500 to 1000 | 1.107 | 0.286 | -0.854 | 0.407 | 0.063 | -1.734 | 0.104 | **0.021** |
| Faces | ID | 100 to 500 | 4.900 | **<0.001** | -4.565 | **<0.001** | **<0.001** | -6.361 | **<0.001** | **<0.001** |
|  |  | 500 to 1000 | 2.474 | 0.022 | -2.161 | 0.042 | **0.010** | -1.401 | 0.176 | **<0.001** |
|  | Expression | 100 to 500 | 3.036 | 0.006 | -3.266 | 0.004 | **0.007** | -3.211 | **0.004** | **<0.001** |
|  |  | 500 to 1000 | 2.059 | 0.052 | -2.619 | 0.016 | **0.002** | -2.169 | **0.042** | **0.001** |

### 2.7 LDA-based decoding methods

To assess whether the results generalized to another decoding algorithm, we repeated the main analyses using LDA instead of SVM. To examine the effect of regularization strength, we performed the decoding procedure on each dataset using a range of values (0.001, 0.01, 0.1, and 1). For these analyses, we used 3-fold cross-validation (*N* = 3), with the maximum number of trials per average (*T*) permitted for that *N* in each dataset. Note that the regularization parameter for LDA, λ, is different from the *C* parameter used in SVM. It varies between 0 and 1, and it modifies the covariance matrix by "shrinking" it toward a simpler structure.

Similar to the SVM analyses, to examine the effects of the cross-validation parameters *N* and *T*, we held the regularization strength constant at a neutral level (λ = 0, which is the default value in MATLAB’s function) and repeated the decoding procedure for each dataset across a range of *N* and *T* values. We used the same range of *N* values as in the SVM analyses, except that *N* = 2 was not included for LDA. This is because 2-fold cross-validation yields training sets containing only one pseudotrial for each class, which can result in unstable estimates of the class means and, critically, the shared covariance matrix required by LDA.

## 3 Results

### 3.1 Binary decoding cases

This section presents the decoding results for the ERP CORE paradigms, each of which involved binary classification. Note that the differences between experimental conditions for these ERP components could be easily detected using conventional univariate analyses (see Kappenman et al., 2021).

The top row of Figure 2 shows how mean decoding accuracy and effect size varied across different regularization settings in the ERP CORE experiments. For most ERP components, both metrics showed very little change as a function of the regularization strength. However, for the N2pc and LRP components, decoding accuracy showed only a modest decrease (approximately 3–5 %), whereas the reduction was more pronounced for effect size when the regularization strength was less than 0.1.

**Figure 2.**
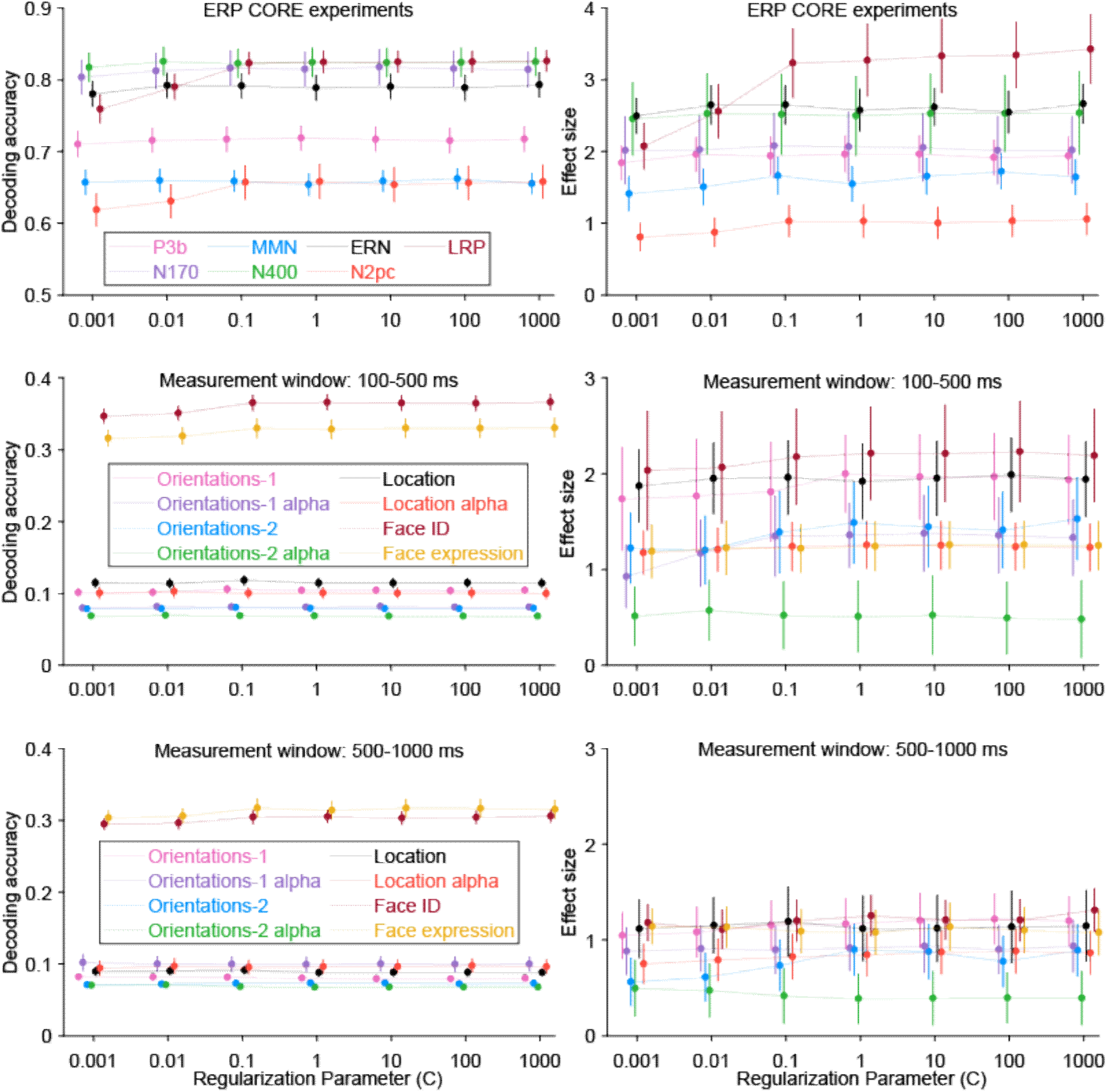
Decoding accuracy (left column) and effect size (right column) for different regularization strengths, separately for each dataset. Error bars show the standard error for each value. The chance level for decoding accuracy is 0.5 for the ERP CORE experiments, 0.25 for the Face experiment (face ID, face expression), and 0.0625 for the Orientation experiments. Chance is always 0 for effect size.

The top row of Figure 3 shows how mean decoding accuracy varied as a function of the number of crossfolds (N) and the number of trials per average (T) for the ERP CORE paradigms. Decoding accuracy improved as T increased from 2 to at least 18 trials for each ERP component. We observed a peak and then a decline in decoding accuracy as T increased for the LRP and N2pc components. The other components appeared to be nearing a peak at the maximum value of T, but we did not have enough trials to determine conclusively whether decoding accuracy would eventually start to fall. In all cases, the highest decoding accuracy was achieved when N was between 2 and 5. This suggests that having clean data (i.e., many trials per average) is more important for maximizing decoding accuracy than having a large number of training cases (i.e., a large number of crossfolds).

**Figure 3.**
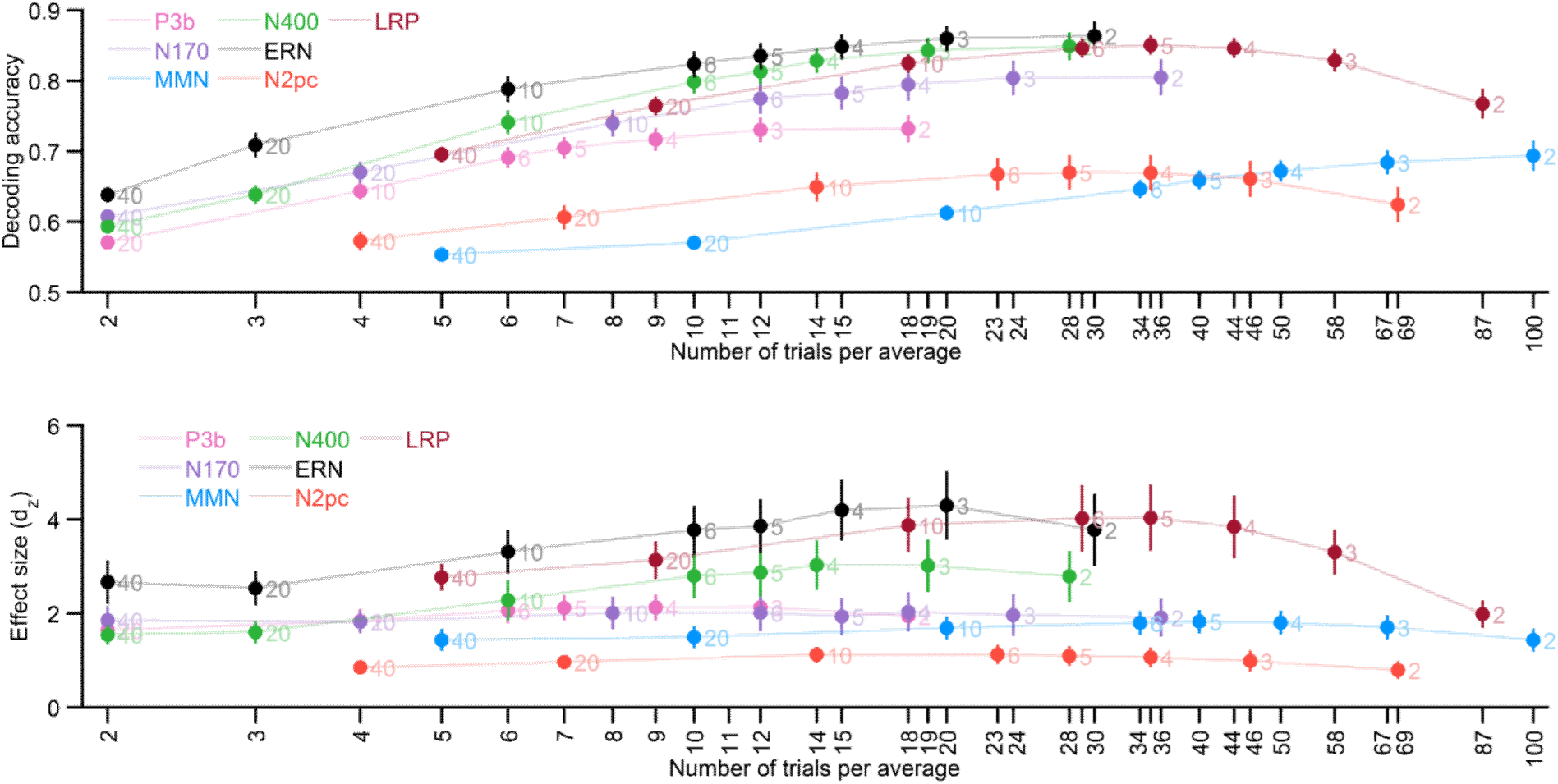
Decoding accuracy (top row) and effect size (bottom row) for different numbers of crossfolds for the ERP CORE paradigms, averaged across the time-window shown in Table 1. The X axis labels indicate the number of trials per average, and the number next to each data point indicates the corresponding number of crossfolds. Error bars show the standard error for each value. The chance level is 0.5 for decoding accuracy and 0.0 for effect size in all cases.

The bottom row of Figure 3 shows the corresponding effect sizes, which take into account both the mean decoding accuracy and the standard deviation across participants. The impact of variations in N and T was not as pronounced for effect size as it was for mean decoding accuracy. In all cases, the effect size increased at least slightly as T increased, then reached a peak and began to decline as T increased further. The optimal T varied quite a bit across components, with the largest effect size occurring between 7 and 12 trials per average for the P3b component and between 34 and 50 trials per average for the MMN. In all cases, the effect size was maximized when N was between 3 and 10. However, variations in N and T had only a minimal impact on the effect size except for the N400, ERN, and LRP components. Note that, whereas the mean decoding accuracy was maximal with only 2 crossfolds in several cases, the maximum effect size required at least 3 crossfolds in all cases. Supplementary Figure S11 shows that the same pattern was observed when the effect size was quantified separately at each time point, and then the effect sizes were averaged across the time window, which better characterizes the central tendency of the single-point effect sizes.

To illustrate the relationship between ERP dynamics and decoding performance, we plotted the ERP difference waves together with the corresponding time-resolved decoding accuracy for the ERP CORE paradigms (Figure 4). In each panel, the ERP difference waveform between the two experimental conditions is shown along with decoding accuracy curves corresponding to classifying between those two conditions, with separate curves representing the different parameter settings (see the left column of Figure 4 for the different *N/T* combinations and the right column for the different *C* values).

**Figure 4.**
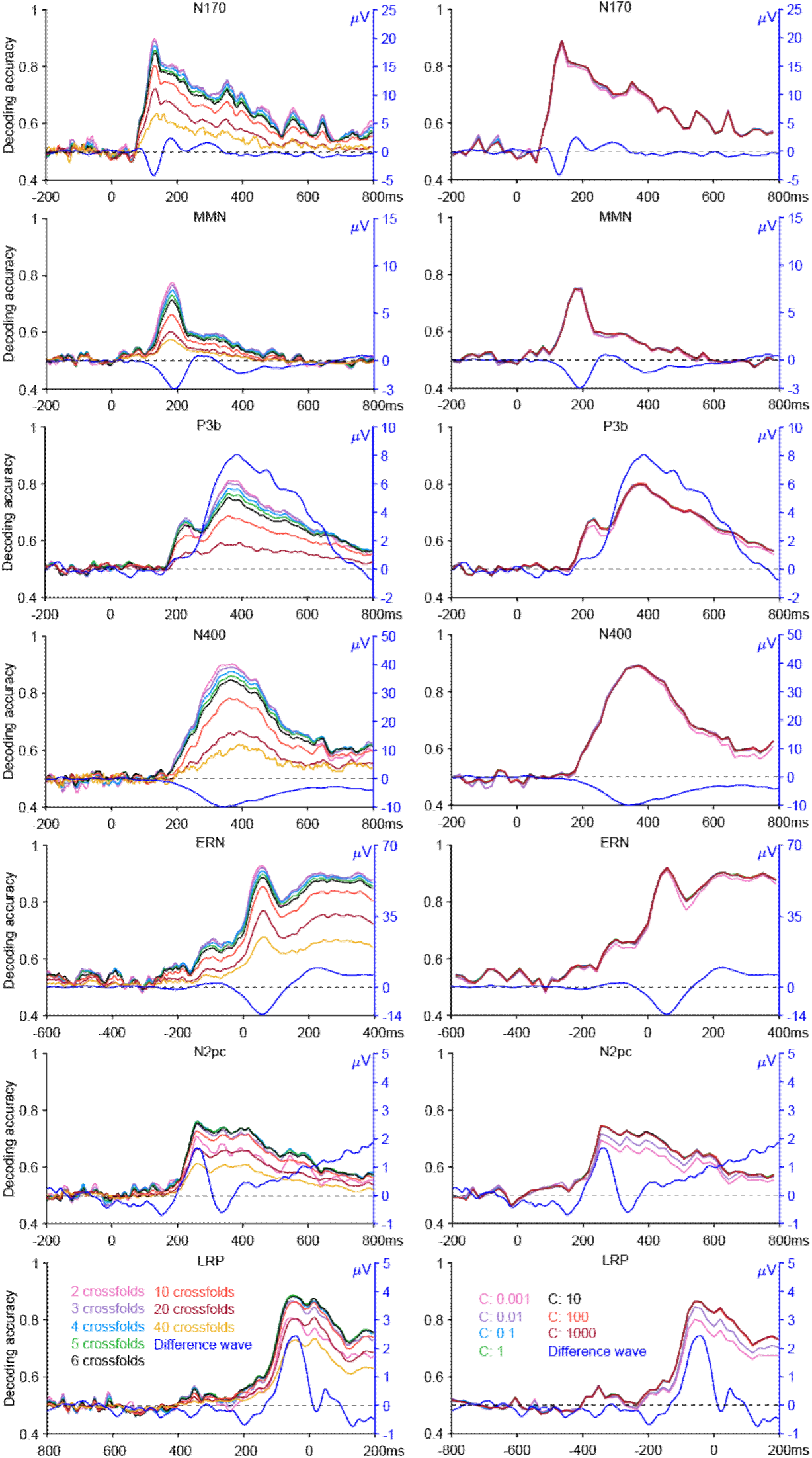
ERP difference waves between experimental conditions (blue curves) and decoding time courses for distinguishing between those same conditions in each of the ERP CORE paradigms. The different decoding curves correspond to different parameter settings (e.g., regularization strength or N/T combinations). The dotted line represents the chance level (0.5) for the decoding time courses and represents zero µV for ERP difference waveforms. The decoding accuracy scale is given along the left side of each panel, and the difference wave amplitude scale is given along the right side.

Across paradigms, decoding accuracy was generally well above chance within the time windows where the corresponding difference wave was clearly different from zero, although in some cases decoding accuracy was well above chance even when the difference wave was close to zero. This likely reflects the fact that decoding is based on all the electrodes, whereas the difference waves show the single “best” electrode for the component of interest. Figure 4 also shows that the effects of the variations in the decoding parameters were similar across the different time points in the decoding waveforms.

Note that decoding accuracy hovered close to the chance value during the prestimulus baseline period in all cases. This is important to verify, because incorrect decoding methods (e.g., failure to equate *T* across classes) will often lead to consistently above-chance or below-chance decoding accuracy during the baseline period. If decoding accuracy is close to chance during the baseline period, this provides some reassurance that the decoding procedure does not contain any biases.

We conducted regression analyses to determine whether the observed differences across regularization parameters and numbers of crossfolds were statistically significant. Because the effects of the number of crossfolds were often nonmonotonic, we included both linear and quadratic terms in the regressions. As shown in Table 3, we found that regularization strength produced significant linear or quadratic effects in only a small subset of cases. However, both the linear and quadratic effects of the number of crossfolds were significant in most cases.

### 3.2 Multiclass decoding cases

We also examined more challenging decoding cases in which the number of classes was greater (4 or 16) and the differences between classes would be difficult to detect using conventional univariate analyses. Both a perceptual time window (100–500 ms) and a working memory time window (500–1000 ms) were analyzed for these paradigms, and we decoded alpha-band power in addition to ERP amplitude in the Orientations paradigms.

The effects of varying the regularization strength for the Orientations and Faces datasets are shown in the middle and bottom rows of Figure 2. Decoding accuracy (left column) showed only slight variations as the regularization strength increased. Effect size (right column) exhibited more noticeable changes in some cases, dropping when the regularization strength was less than 1.

The effect of the cross-validation parameters on decoding accuracy is shown in the top row of Figure 5 for the 100-500 ms time window and in the top row of Figure 6 for the 500-1000 ms time window. Decoding accuracy increased as T increased up to 20-60 trials per average (corresponding to N = 2-5 crossfolds) and then began to decline. As observed for the ERP CORE paradigms, variations in N and T had a smaller impact on the effect size than on the mean decoding accuracy (bottom row in Figures 4 and 5), and the peak effect size was observed with a smaller T and larger N for effect size than for mean decoding accuracy. Specifically, the maximum effect size occurred with a T of 5–32 trials per average and an N of 5-20 crossfolds for most cases. Supplementary Figure S11 shows that the same pattern was observed when the effect size was quantified separately at each time point and then the effect sizes were averaged across the time window.

**Figure 5.**
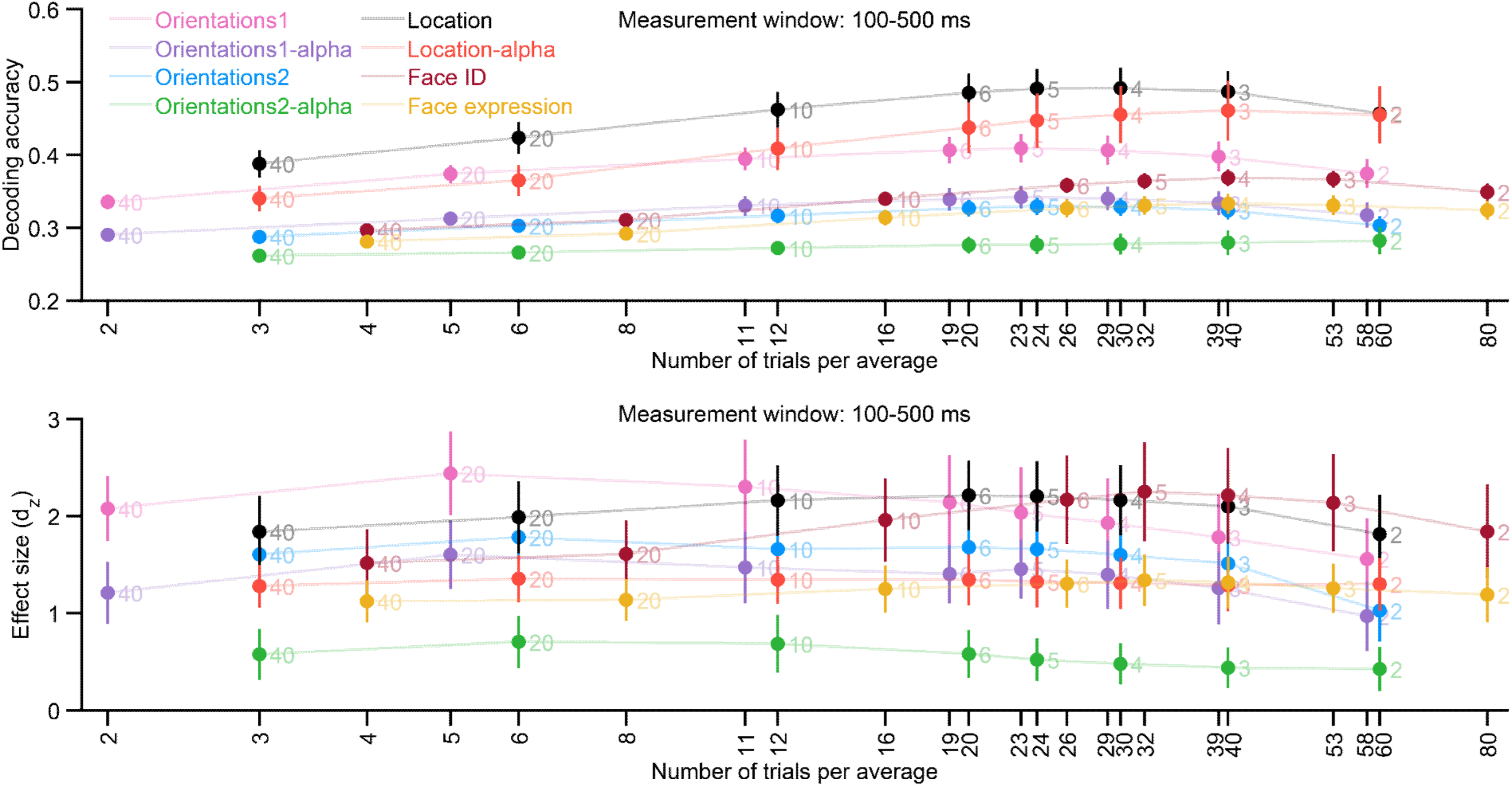
Decoding accuracy (top row) and effect size (bottom row) for different numbers of crossfolds when decoding orientation in the Orientations-1 & 2 datasets when decoding location of the Orientations-2 dataset, and when decoding face identity or facial expression in the Faces dataset. Decoding accuracy was averaged across the perceptual time-window of 100-500 ms. The X axis labels indicate the number of trials per average, and the number next to each data point indicates the corresponding number of crossfolds. Error bars show the standard error for each value. The chance level is 0.25 for decoding accuracy and 0.0 for effect size in all cases.

**Figure 6.**
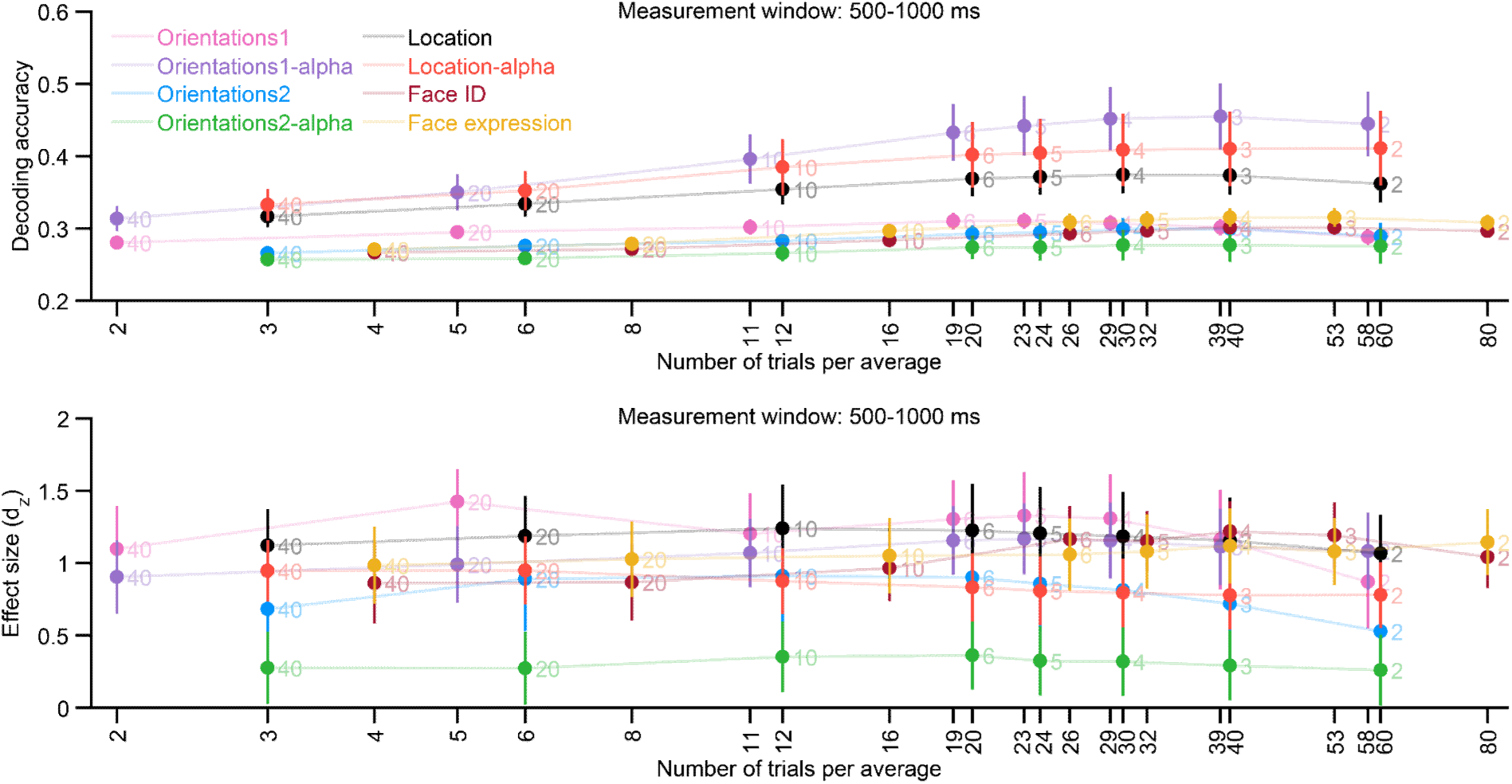
Decoding accuracy (top row) and effect size (bottom row) for different numbers of crossfolds when decoding orientation in the Orientations-1 & 2 datasets when decoding location of the Orientations-2 dataset, and when decoding face identity or facial expression in the Faces dataset. Decoding accuracy was averaged across the working memory time-window of 500-1000 ms. The X axis labels indicate the number of trials per average, and the number next to each data point indicates the corresponding number of crossfolds. Error bars show the standard error for each value. The chance level is 0.25 for decoding accuracy and 0.0 for effect size in all cases.

As shown in Table 3, we found that regularization strength had significant linear or quadratic effects in only a small subset of cases. However, both the linear and quadratic effects of the number of crossfolds were significant in most cases.

### 3.3 Generalization to LDA

We repeated the above analyses using LDA in place of SVM (and varying the LDA λ regularization parameter in place of the SVM *C* parameter). When LDA was used for decoding, varying the λ regularization parameter had minimal impact in any of the datasets except for a modest drop in decoding accuracy and effect size for a λ of 1 in some cases (see supplementary Figure S5). The effects of varying the cross-validation parameters for LDA were similar to the effects observed for SVM with the ERP CORE datasets, except that decoding performance for LDA tended to peak at 4-6 folds instead of 3-5 folds (see supplementary Figure S6). For the multiclass decoding cases, decoding performance tended to peak at 3-5 folds for LDA just as for SVM.

## 4 Discussion

This study investigated how regularization and cross-validation parameters influence EEG/ERP decoding performance. To our knowledge, this is the first study to systematically explore these effects across a wide range of EEG/ERP paradigms involving different electrode configurations, sample sizes, and experimental conditions. Using the ERP CORE, Orientations-1, Orientations-2, and Faces datasets, we found that regularization strengths of less than C = 1 negatively impacted mean decoding accuracy and the corresponding effect size compared to a strength of C = 1, with little or no impact of increasing the regularization strength above 1. The analyses of the cross-validation parameters indicated that keeping the number of crossfolds low (N = 2-5) and the number of trials per average correspondingly high (typically T = 10-50) yielded the highest mean decoding accuracy. By contrast, the effect size (which takes into account both the mean and the standard deviation across participants) benefited from more crossfolds (typically N = 3-10) even though this meant fewer trials per average (typically T = 5-30). Note, however, that the effects of variations in N and T on the effect size were modest in most cases, especially when N and T were not extreme.

### 4.1 Effect of regularization strength

The present findings provide quantitative evidence about the optimal regularization strength, whereas previous work has often relied on default or heuristic settings without systematic evaluation (Blankertz et al., 2011; Grootswagers et al., 2017). Regularization in SVM controls the trade-off between minimizing classification errors during training and improving generalization to the test data by maximizing the margin (Cortes & Vapnik, 1995), with values of C less than 1 favoring generalization. Interestingly, we found that decoding accuracy for the test set (and the corresponding effect size) declined when C was substantially less than 1 (when the regularization was very strong, with very wide margins). This suggests that high levels of regularization (C << 1) did not allow sufficiently good fitting of the training data, which then led to poor results for the test data. However, we never saw a substantial benefit of having C exceed 1, so we recommend using a value of 1, which provides an equal balance between good fitting of the training data and good generalization to the test data. However, this recommendation may not generalize to datasets that are quite different from those examined here. For example, it may not generalize to data obtained with small numbers of electrodes, noisier recording setups (e.g., dry electrodes; G.-L. Li et al., 2020), or very different participant populations (e.g., infants; Ashton et al., 2022; Turk-Browne & Aslin, 2024).

### 4.2 Effect of the cross-validation parameters

Increasing the number of crossfolds may be beneficial for decoding by increasing the number of training cases, but it may be detrimental by decreasing the number of trials per average and therefore the SNR of both the training data and the test data. Consistent with these competing factors, we found that decoding was typically best when extreme values of T were avoided and the number of crossfolds/trials was moderate to low. However, the optimal balance was somewhat different depending on whether we examined mean decoding accuracy or the effect size. Mean decoding accuracy typically benefited from maximizing the number of trials per average, as long as the number of crossfolds did not get too small (e.g., N = 2). By contrast, effect size benefited from increasing the number of crossfolds somewhat (e.g., N = 5-10) at the cost of fewer trials per average. Importantly, increases in decoding accuracy obtained with small numbers of crossfolds and large numbers of trials per average may also be accompanied by increased variability across participants or iterations. As a result, decoding accuracy values that appear to exceed the theoretical chance level may still fall within a relatively broad confidence interval around chance performance. Consequently, relying solely on mean decoding accuracy may sometimes lead to overinterpretation of small effects. The present study partly addresses this issue by also examining effect sizes, which take variability into account in addition to mean decoding performance. Maximizing the decoding accuracy is primarily important in engineering applications, whereas maximizing the effect size (and therefore statistical power) is usually more important in scientific studies (see Hebart & Baker, 2018). Most scientific studies should therefore avoid low numbers of crossfolds. Note, however, that the impact of the number of trials/crossfolds on effect size was relatively modest in most cases. As a result, researchers should be able to obtain good effect sizes as long as they avoid extreme numbers of trials or crossfolds. For experiments like those examined here, researchers should be able to achieve near maximum performance for both decoding accuracy and effect size by using N = 3-5 crossfolds with at least T = 10 trials per average. Although one might wish to define a single recommended value, some combinations of N and T may be more natural than others for a given dataset (e.g., N=3 and T=15 when the total number of trials is 45 per class). Therefore, the ranges identified here should be viewed as approximate guidelines rather than strict rules.

Because parameters such as the number of crossfolds (N) and the number of trials per average (T) can influence decoding performance, it may be tempting to explore a large number of parameter combinations and select the configuration that yields the highest decoding accuracy for a given dataset. However, this type of analytic flexibility may increase the risk of inflated effect estimates or false positives, similar to other forms of researcher degrees of freedom in data analysis. Therefore, when decoding analyses are used in a hypothesis-testing framework, it is generally advisable to determine these parameters *a priori*—based on prior literature, methodological considerations, or independent datasets—rather than selecting them based on the results obtained in the dataset being analyzed. One goal of the present study was to provide empirical guidance about how decoding results vary across commonly used parameter ranges, which may help researchers make a priori parameter choices and reduce analytic flexibility.

When pseudotrials are constructed by averaging together relatively large numbers of trials, the resulting signals resemble high signal-to-noise ERPs. In such cases, decoding performance will be related to the same underlying condition differences that would also be observable in conventional ERP analyses. Consequently, decoding approaches using small N and large T values may become conceptually closer to comparing ERPs. Nevertheless, multivariate decoding differs from traditional ERP analyses in that it evaluates whether distributed patterns of activity across electrodes contain information that discriminates experimental conditions, rather than focusing on amplitude differences at predefined electrodes and time windows. Thus, decoding can reveal spatially distributed patterns that may be individually weak at single electrodes but jointly informative across the electrode array. In addition, decoding is performed separately for each subject, which allows for the possibility of individual differences in the patterns that discriminate among conditions.

Conventional univariate analyses, in contrast, look for commonalities across subjects in the topographic pattern of differences among conditions, which may be insensitive when individual differences are present. Because of these differences, decoding may often be more sensitive to differences among conditions than conventional ERP analyses. Indeed, Carrasco et al. (2024) demonstrated that effect sizes are often increased when decoding is used instead of conventional univariate ERP analyses to quantify differences among conditions. Thus, pseudotrial-based decoding should be viewed as complementary to conventional ERP analyses.

The decoding results were fairly consistent across a wide range of experimental paradigms, which varied in sample size, the electrode densities, the nature of the experimental conditions, and the number of classes. Consequently, the conclusion that decoding performance benefits from using N = 3-5 and T >= 10 will likely generalize to a broad range of datasets. Note that, because we were focused on typical cases in which the total number of trials is fixed and N and T are therefore reciprocally related, we did not examine the effects of variations in T for a fixed value of N or variations in N for a fixed value of T. However, Scrivener et al. (2023) demonstrated with simulated EEG data that increasing T improved decoding accuracy when N was held constant. Presumably, increasing N while holding T constant would also increase decoding accuracy by increasing the amount of training data.

Note that the benefit of increasing T varied somewhat across datasets (see Figure 3). One possible (but untested) explanation is that ERPs with higher within-condition trial variance or weaker between-condition separability may benefit more from the noise reduction produced by averaging, whereas ERPs with more stable and well-separated responses may show smaller benefits of averaging. However, because the present study was not designed to quantify these variance components directly, this interpretation remains tentative. A systematic investigation linking ERP-specific signal characteristics to decoding performance under different averaging strategies will be an important direction for future research.

Interestingly, restricting the signal to the alpha band reduced both decoding accuracy and effect size for some cases (see Figure 5), even in paradigms where alpha oscillations are thought to be functionally relevant. It is possible that decoding performance for broadband ERP signals benefits from access to the full spatiotemporal signal structure, whereas restricting the signal to the alpha band reduces the dimensional richness of the data available to the classifier. However, because the present study was not designed to explore differences in decoding accuracy between oscillatory signals and phase-locked broadband signals, this is only speculation, and the finding of smaller effect sizes for alpha-band decoding should be treated as tentative.

### 4.3 Generalizability

Although the present results provide practical guidance for selecting decoding parameters, these recommendations should not be interpreted as rigid rules. For datasets with characteristics similar to those examined here (e.g., typical ERP paradigms with adult participants and moderate numbers of trials), the suggested parameter ranges may serve as a reasonable starting point and justification for analysis choices, especially given the fair amount of consistency we observed across datasets. For very different studies, however, researchers may benefit from collecting pilot data or evaluating parameter choices on independent datasets to determine how different parameter combinations influence decoding performance in their specific task. In particular, all of the datasets examined here were collected from similar adult populations (i.e., highly cooperative college students), using similar high-quality gel-based recording systems and analyzed through comparable processing pipelines.

Because we did not include other types of datasets collected from different populations, such as infants, children, and people with brain disorders (Marsicano et al., 2024; Ng et al., 2022; Turoman et al., 2024), from different recording systems (e.g., dry electrodes Habibzadeh Tonekabony Shad et al., 2020; G.-L. Li et al., 2020), from mobile EEG systems (Niso et al., 2023; Park et al., 2015), or from different recording environments (e.g., hospitals; Lei et al., 2024), we simply cannot know whether our findings would generalize to those contexts. Additional research would be needed to determine the optimal averaging and regularization parameters for these contexts.

Additionally, although some previous studies have shown that single-trial classification can yield similar results to the averaged-trial approach (Williams et al., 2020), it is unknown whether the present conclusions about regularization will generalize to single-trial classification. This question was beyond the scope of the current study.

We repeated the main analyses with LDA in place of SVM. We found very little impact of the LDA λ regularization parameter as long as the λ value was less than 1. When we examined the cross-validation parameters with LDA, the pattern of results was much like that obtained with SVM, except that decoding performance benefited from slightly more crossfolds with LDA than with SVM (4-6 vs. 3-5) with the ERP CORE datasets. SVM may be able to tolerate fewer crossfolds (and therefore fewer training cases) because it focuses on cases near the decision hyperplane, whereas LDA takes into account the entire distribution of training values. However, the difference between LDA and SVM was modest, and 3-5 crossfolds was found to be optimal for both methods in the multiclass decoding datasets. Thus, our recommendations for LDA are similar to those for SVM, except that LDA uses a different regularization parameter for which we have a different recommendation (0.1 or less), and we recommend 4-6 crossfolds when LDA is used to perform binary decoding.

It is also unknown whether the present results will also generalize to other relatively simple classification algorithms, such as the random forest approach (Breiman, 2001), and to more advanced algorithms, such as Bayesian classification, convolutional neural networks, recurrent neural networks, and deep belief networks (Al-Saegh et al., 2021; Craik et al., 2019; Gong et al., 2022; Hosseini et al., 2021).These approaches may benefit from a different balance between training accuracy and test accuracy (i.e., regularization strength) or a different balance between the number of classes and the number of trials per class. This is an issue that deserves study, especially in cases where these other approaches are superior to SVMs. However, the SVM and LDA approaches are very widely used for EEG/ERP decoding at present, so the present conclusions will be valuable to many researchers.

The current SVM analyses focused on linear SVMs, which are commonly used in basic science ERP decoding studies such as those analyzed here because of their robustness and interpretability in high-dimensional neural data. Another option is to use a radial basis function kernel, but this involves multiple hyperparameters (C and γ). We conducted preliminary analyses of the present datasets with Matlab’s default value of γ, and we found poor decoding accuracy.

However, decoding might be better with different γ values, and exploring the entire space of C and γ values in combination with the cross-validation parameters is beyond the scope of the present study. It is impossible to know whether the present results obtained with a linear SVM kernel would generalize to nonlinear SVM kernels, and this issue would need to be assessed with additional research.

In all of the datasets assessed here, averaging was performed across many trials with identical or nearly identical stimuli. Consequently, the present recommendations were derived primarily from conventional ERP paradigms in which pseudotrial averaging mainly serves to reduce trial-level noise. An important open question is whether these conclusions generalize to decoding paradigms that use large numbers of trial-unique exemplars within each category, with little or no repetition of the individual exemplars. In these cases, the goal of including many trials is typically to assess exemplar-independent representations of stimulus categories, not simply to increase the signal-to-noise ratio. To assess whether the patterns observed here generalize to such cases, we conducted an additional analysis on a large-scale categorization dataset in which many different exemplars were presented from the categories of birds, vehicles, mammals, and fish (Poncet et al., 2025). As observed in our other analyses, variations in the SVM regularization parameter had little or no impact on decoding performance (see supplementary Figure S8). When we varied the cross-validation parameters, however, we found a somewhat different pattern in this dataset. Specifically, whereas decoding accuracy tended to peak with 3-5 crossfolds in the main set of analyses, decoding accuracy in the Poncet et al. dataset rose monotonically as the number of crossfolds fell, with the peak decoding accuracy observed with 2 crossfolds (see supplementary Figure S9). However, the influence of the number of crossfolds on the *dz* metric of effect size was modest, just as we observed in many of the other datasets.

Because the Poncet et al. (2025) dataset differed in many ways from the other datasets analyzed in the present study, we cannot be certain that the somewhat different pattern observed for decoding accuracy with this dataset was a result of the fact that many different exemplars were being averaged together for each pseudotrial. However, this result does suggest that the present recommendations about the number of crossfolds may not apply to such cases. If the key factor is indeed the fact that many different exemplars were being averaged together, we speculate that the differences in the neural responses produced by the different exemplars are a source of variance that tends to decrease decoding accuracy, and averaging many trials together is especially important in this case to minimize that source of variance. In such paradigms, pseudotrial averaging may serve a somewhat different function than in conventional ERP experiments. Rather than primarily reducing random trial-level noise from repeated presentations of nearly identical stimuli, averaging may also help reduce variability associated with exemplar-specific neural responses while preserving category-level information shared across exemplars. This qualitative difference may alter the optimal balance between the number of crossfolds and the number of trials per average.

Some paradigms included in the present study (e.g., MMN, P3b, and ERN) inherently involve unequal numbers of trials across conditions, which could potentially lead to differences in SNR when trials are grouped into pseudotrials for cross-validation. To minimize this issue, we equated the number of trials across conditions by randomly subsampling the majority condition to match the trial count of the minority condition before constructing pseudotrials. This ensured that each pseudotrial was formed from comparable numbers of trials across conditions.

Nevertheless, class imbalance and associated SNR differences remain an important methodological consideration for decoding analyses, and future work should further investigate strategies for addressing these issues.

### CRediT authorship contribution statement

**Guanghui Zhang**: Writing – review & editing, Writing – original draft, Visualization, Validation, Supervision, Software, Resources, Methodology, Investigation, Project administration, Conceptualization. **Xinran Wang**: Visualization, Validation, Supervision, Software, Resources, Methodology**. Kurt Winsler**: Writing – review & editing, Validation, Visualization, Software. **Steven J. Luck**: Writing – review & editing, Writing – original draft, Visualization, Validation, Supervision, Software, Project administration, Methodology, Funding acquisition, Conceptualization.

## Acknowledgments

We thank the members of the Luck Lab for their valuable contributions to the community and thoughtful insights, including but not limited to: Carlos Daniel Carrasco, John Kiat, David Garrett, and Brett Bahle. We also thank two anonymous reviewers, who suggested several helpful additional analyses.

## Supplementary Materials

### S1. Effects of the cross-validation parameters on decoding performance

**Figure S1.**
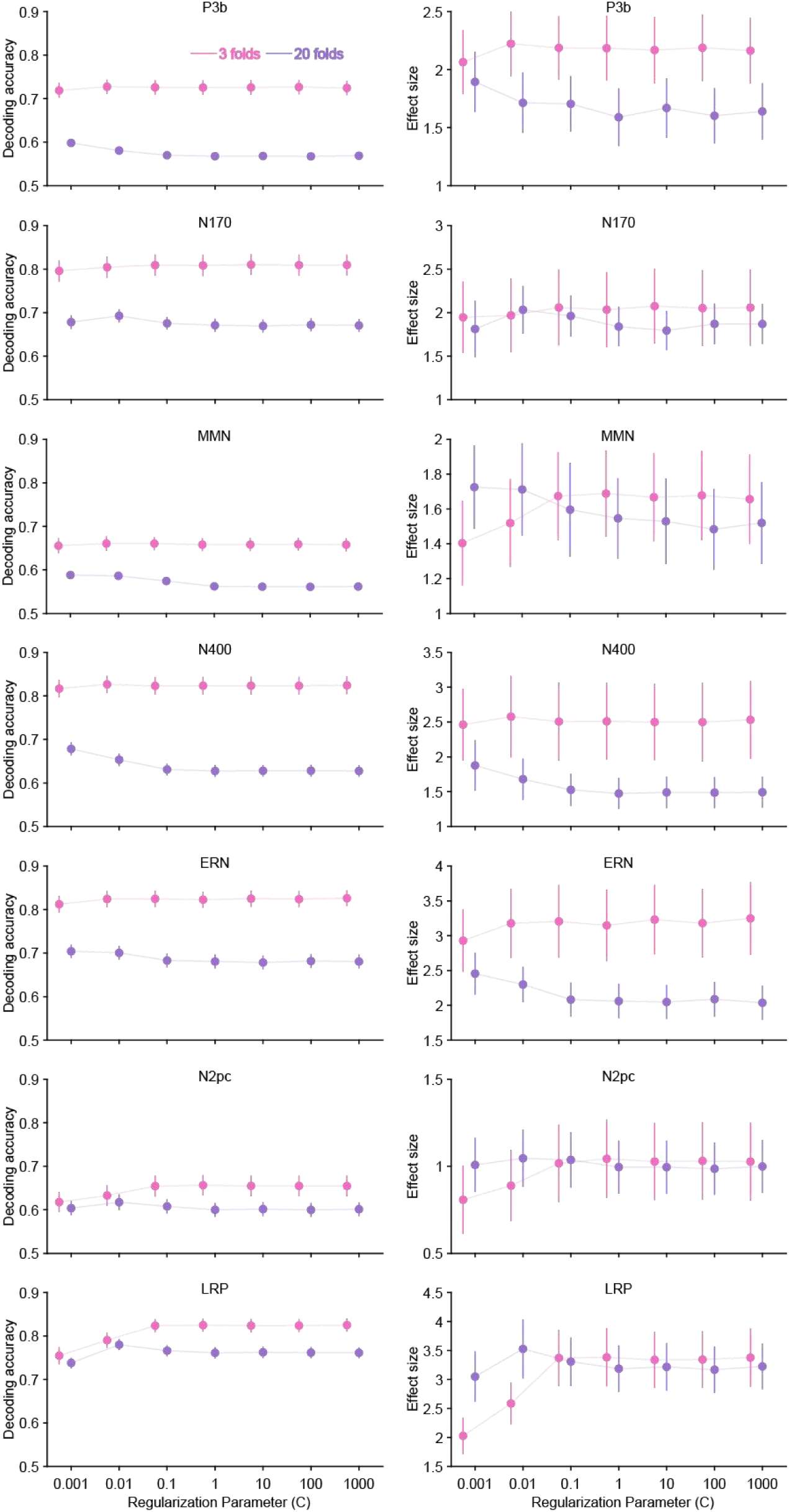
Decoding accuracy (left column) and effect size (right column) for different regularization strengths for ERP CORE dataset for different crossfold numbers (3 vs. 20). Error bars show the standard error for each value.

**Figure S2.**
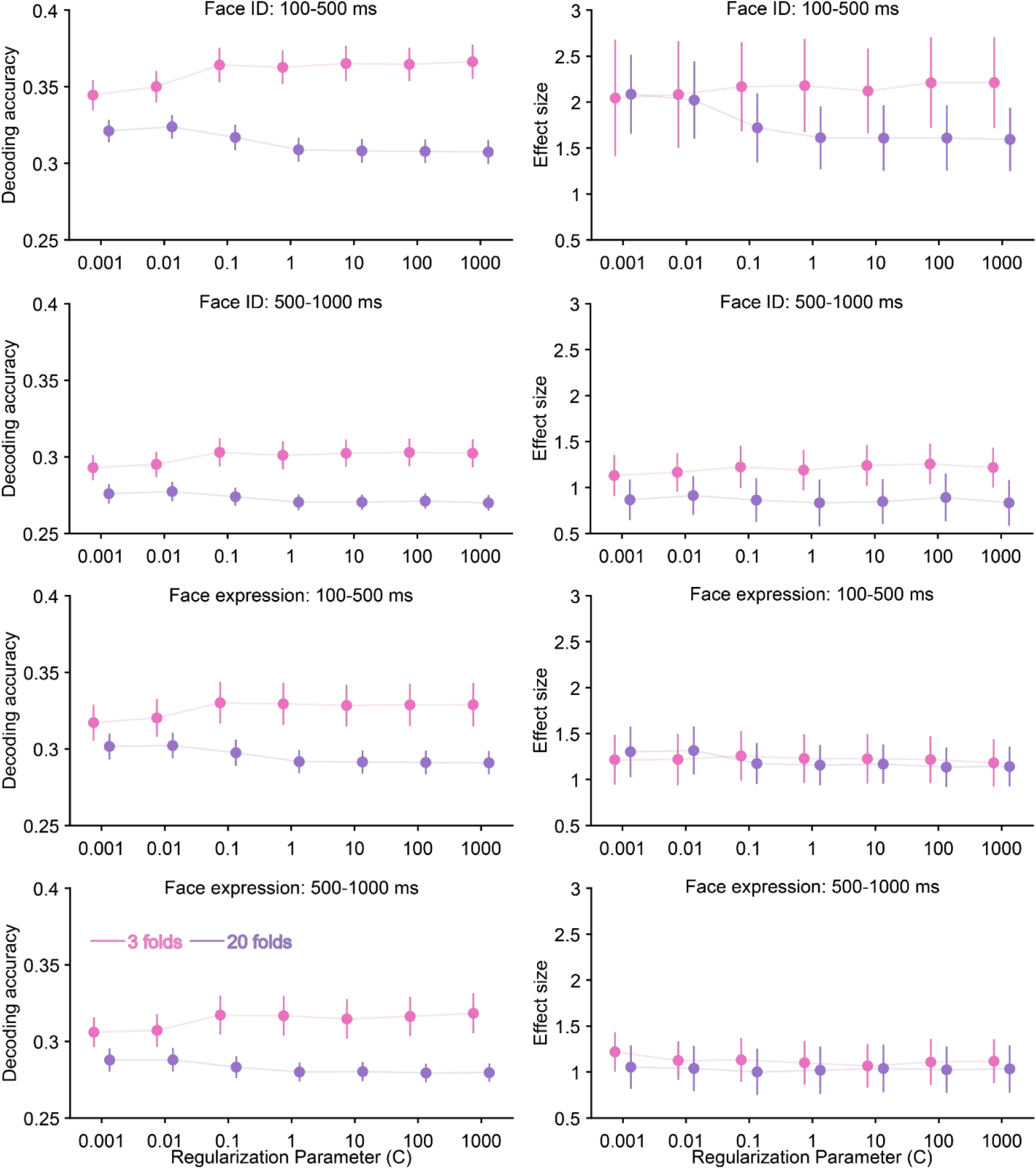
Decoding accuracy (left column) and effect size (right column) for different regularization strengths for the Faces dataset for different crossfold numbers (3 vs. 20). Error bars show the standard error for each value.

**Figure S3.**
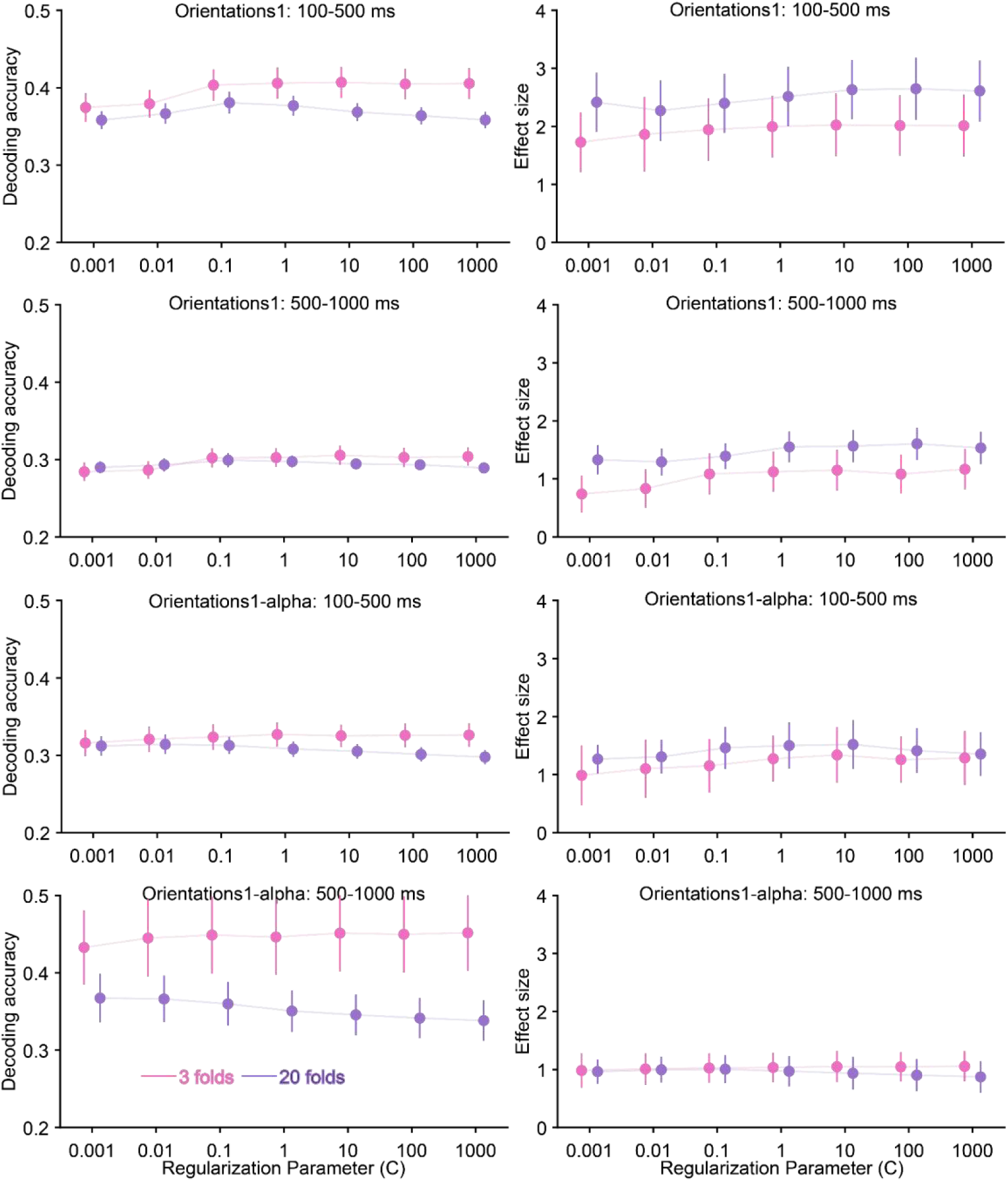
Decoding accuracy (left column) and effect size (right column) for different regularization strengths for the Orientations-1 dataset for different crossfold numbers (3 vs. 20). Error bars show the standard error for each value.

**Figure S4.**
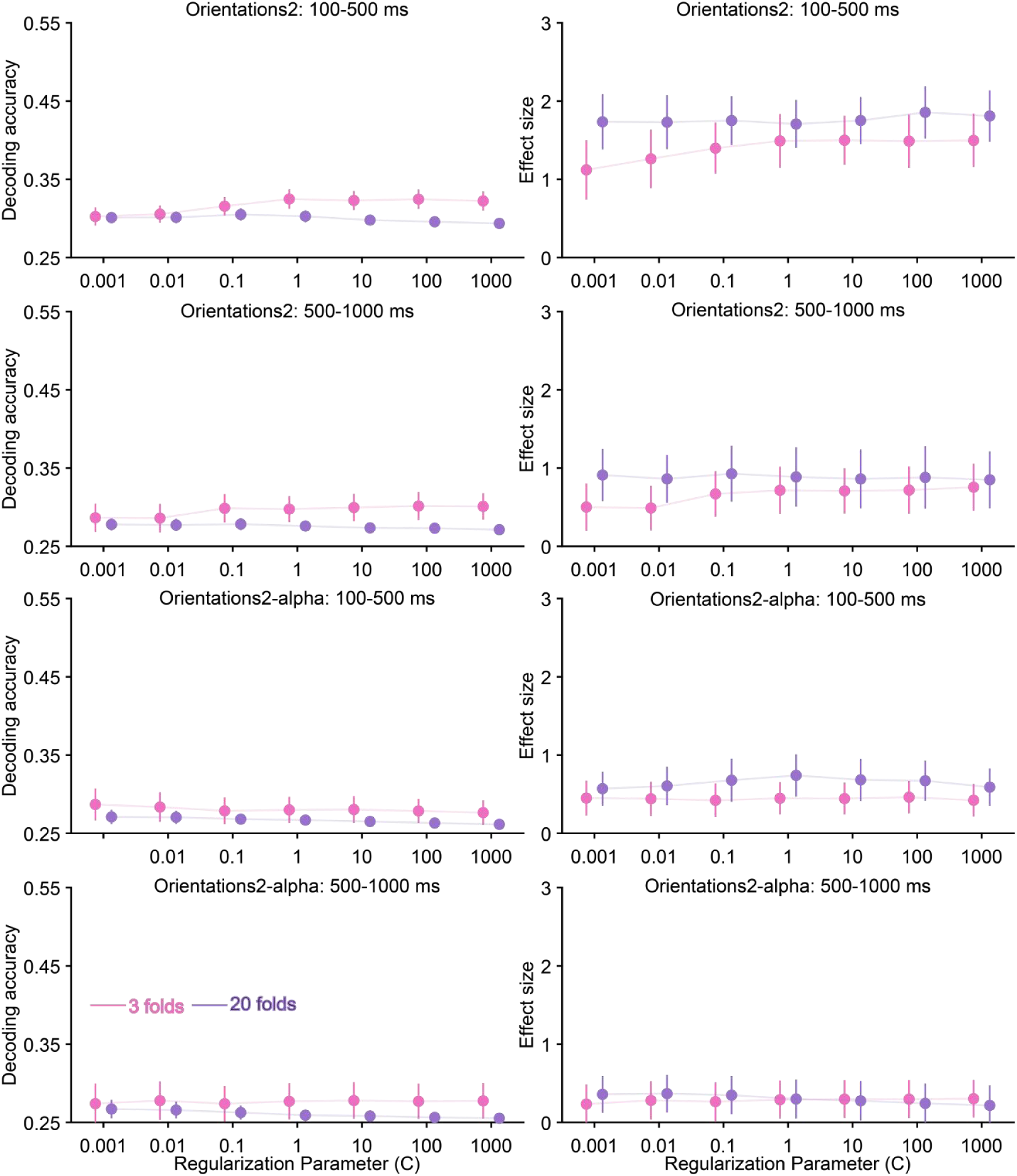
Decoding accuracy (left column) and effect size (right column) for different regularization strengths for the Orientations-2 dataset for different crossfold numbers (3 vs. 20). Error bars show the standard error for each value.

### S2. Effects of the cross-validation parameters on LDA decoding performance

**Figure S5.**
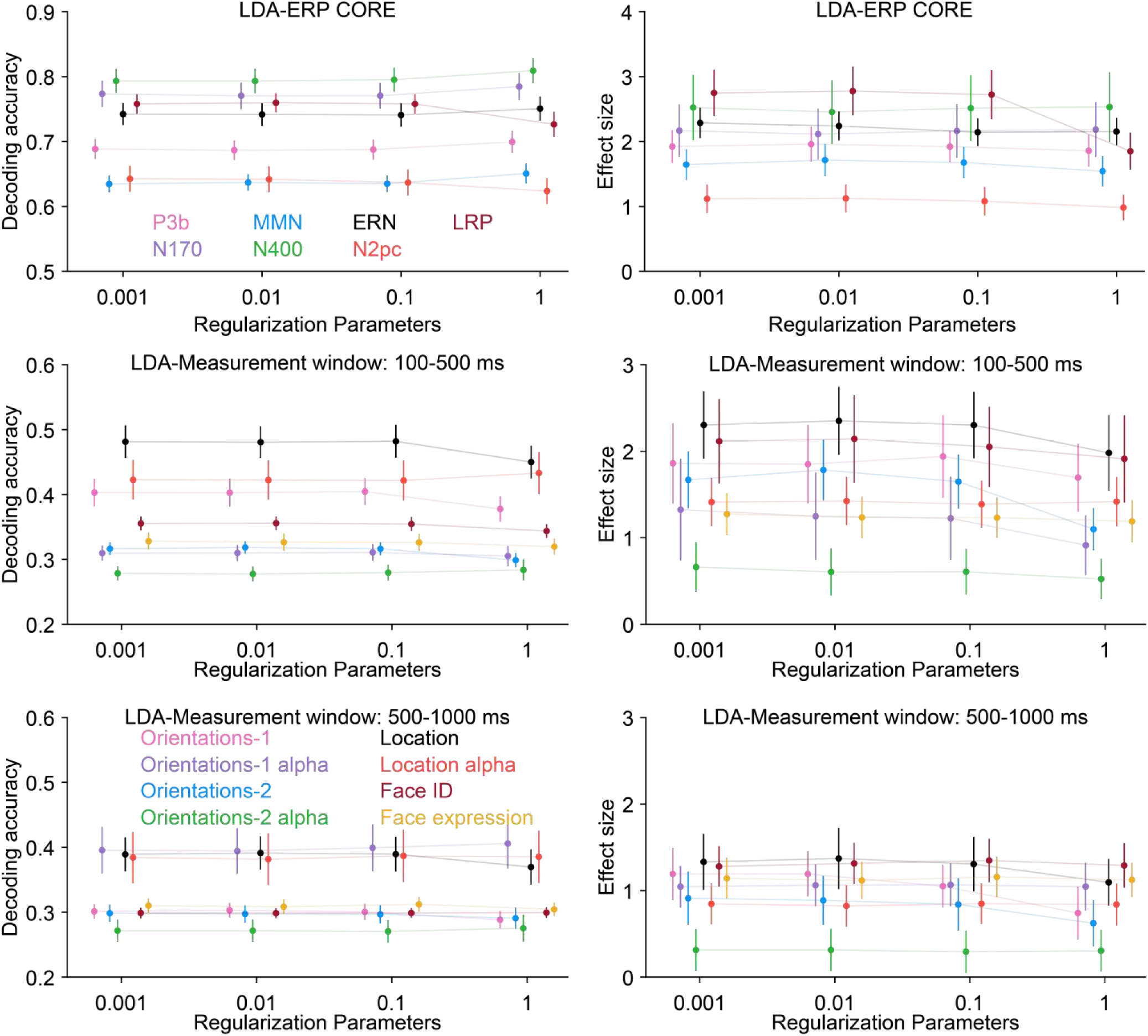
LDA-based decoding accuracy (left column) and effect size (right column) for different regularization parameters, separately for each dataset. Error bars show the standard error for each value. The chance level for decoding accuracy is 0.5 for the ERP CORE experiments, 0.25 for the Face experiment, and 0.25 for the Orientation experiments. The chance level for effect size is zero in all cases.

**Figure S6.**
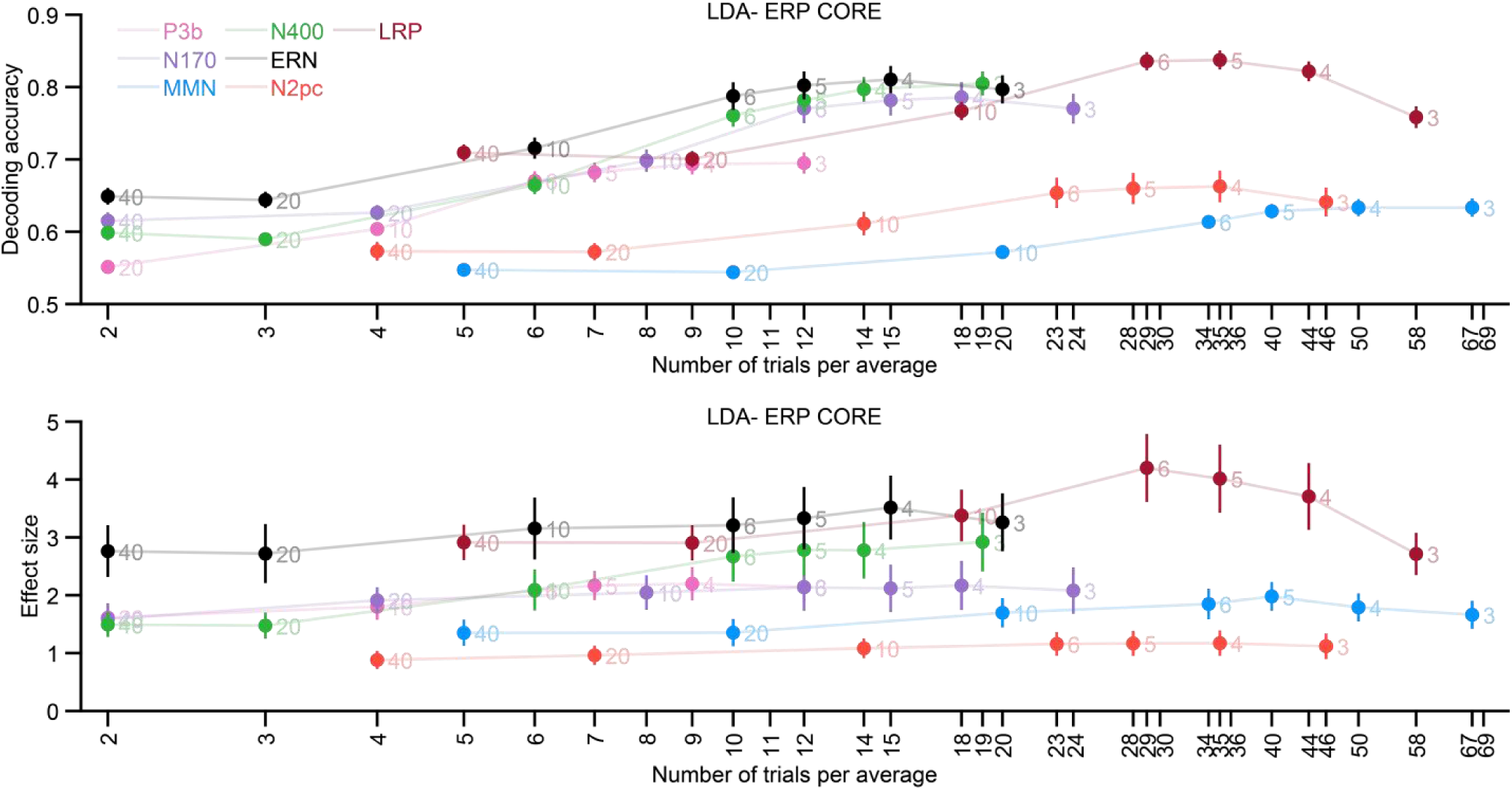
LDA-based decoding accuracy (top row) and effect size (bottom row) for different numbers of crossfolds for the ERP CORE paradigms, averaged across the time-window shown in Table 1. The X axis labels indicate the number of trials per average, and the number next to each data point indicates the corresponding number of crossfolds. Error bars show the standard error for each value. The chance level is 0.5 for decoding accuracy and zero for effect size in all cases.

**Figure S7.**
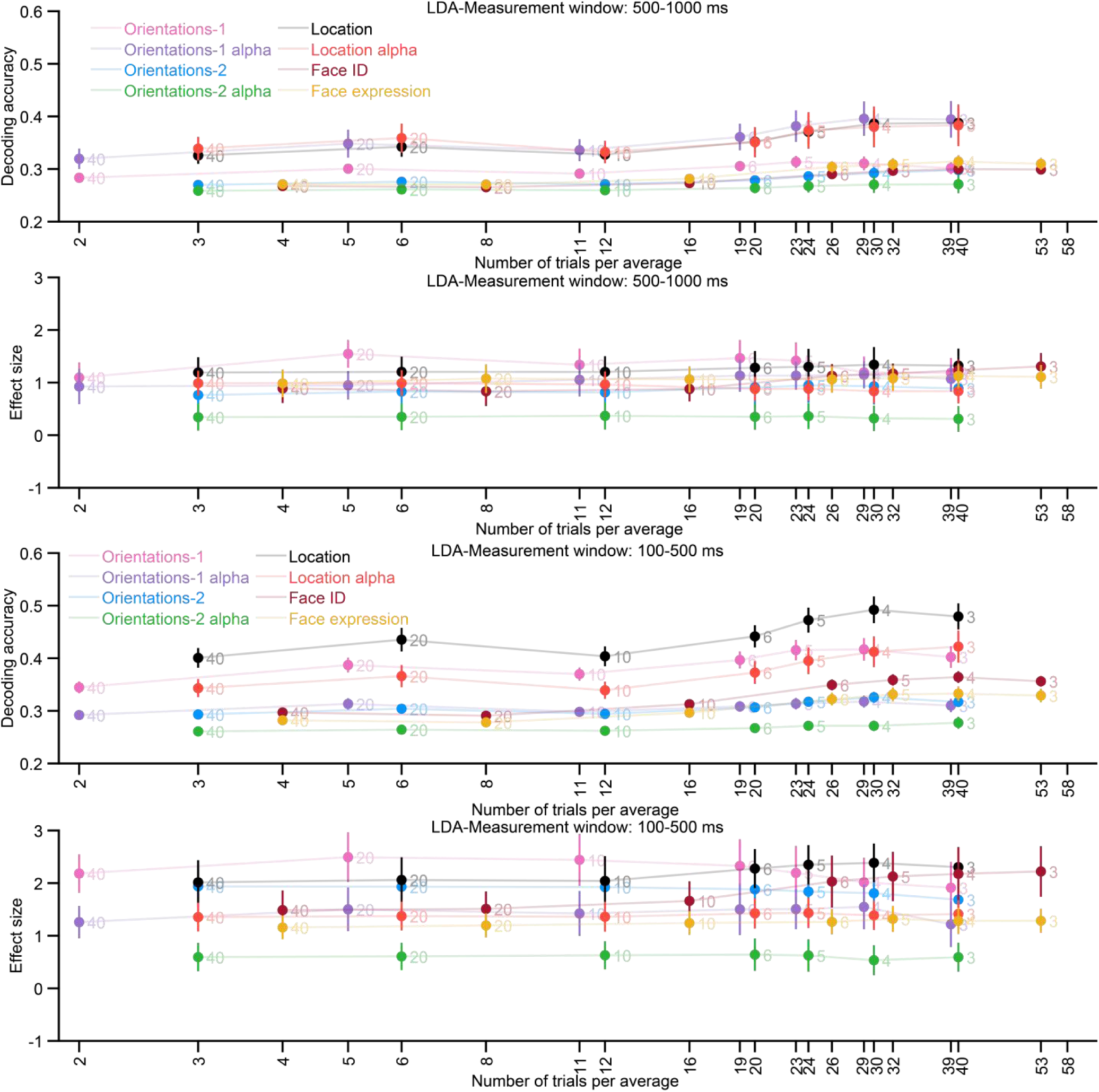
LDA-based decoding accuracy (first and third rows) and effect size (second and fourth rows) for different numbers of crossfolds when decoding orientation in the Orientations-1 & 2 datasets, when decoding location of the Orientations-2 dataset, and when decoding face identity or facial expression in the Faces dataset. Decoding accuracy was averaged across the perceptual time-window of 100-500 ms (the top two rows) and the working memory time-window of 500-1000 ms (the bottom two rows). The X axis labels indicate the number of trials per average, and the number next to each data point indicates the corresponding number of crossfolds. Error bars show the standard error for each value. The chance level is 0.25 for decoding accuracy and zero for effect size in all cases.

### S3. S3SVM-based decoding performance for the paradigm with non-repeated stimulus

**Figure S8.**
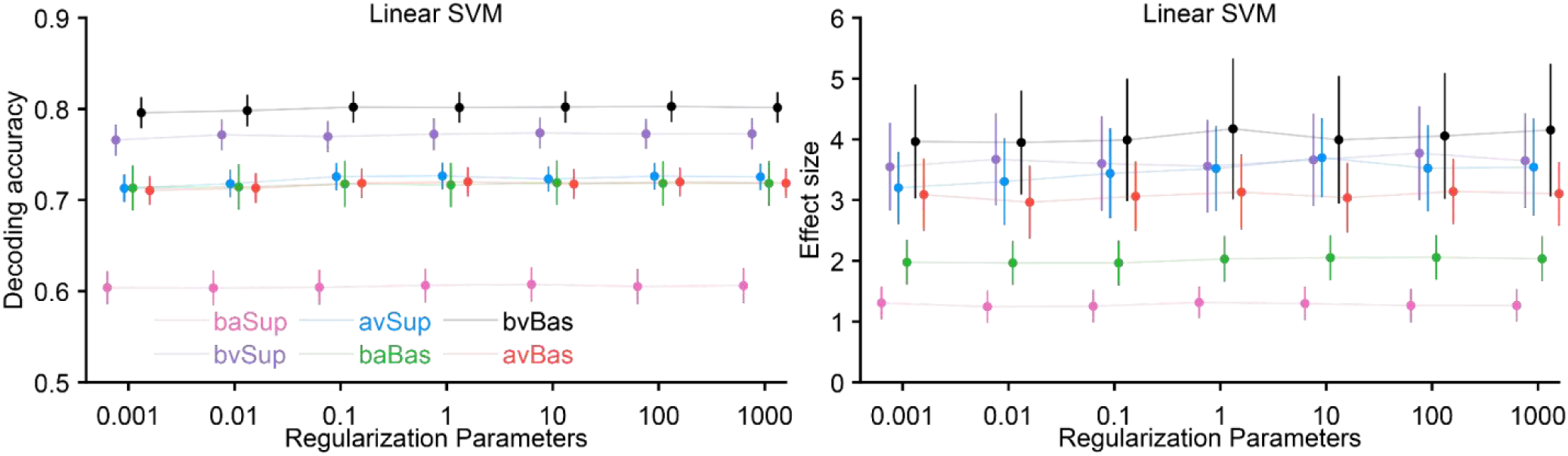
Decoding accuracy (left column) and effect size (right column) for different regularization strengths for the dataset from Poncet et al (2025). The decoding was computed within the time window after stimulus onset (from 100 to 500ms). Error bars show the standard error for each value. The chance level is 0.5 for decoding accuracy and zero for effect size. baSup: Bird/Non-bird animal at the superordinate level; avSup: Non-bird animal/Vehicle at the superordinate level. bvSup: Bird/Vehicle at the superordinate level; baBas: Bird/Non-bird animal at the basic level; avBas: Non-bird animal/Vehicle at the basic level. bvBas: Bird/Vehicle at the basic level.

**Figure S9.**
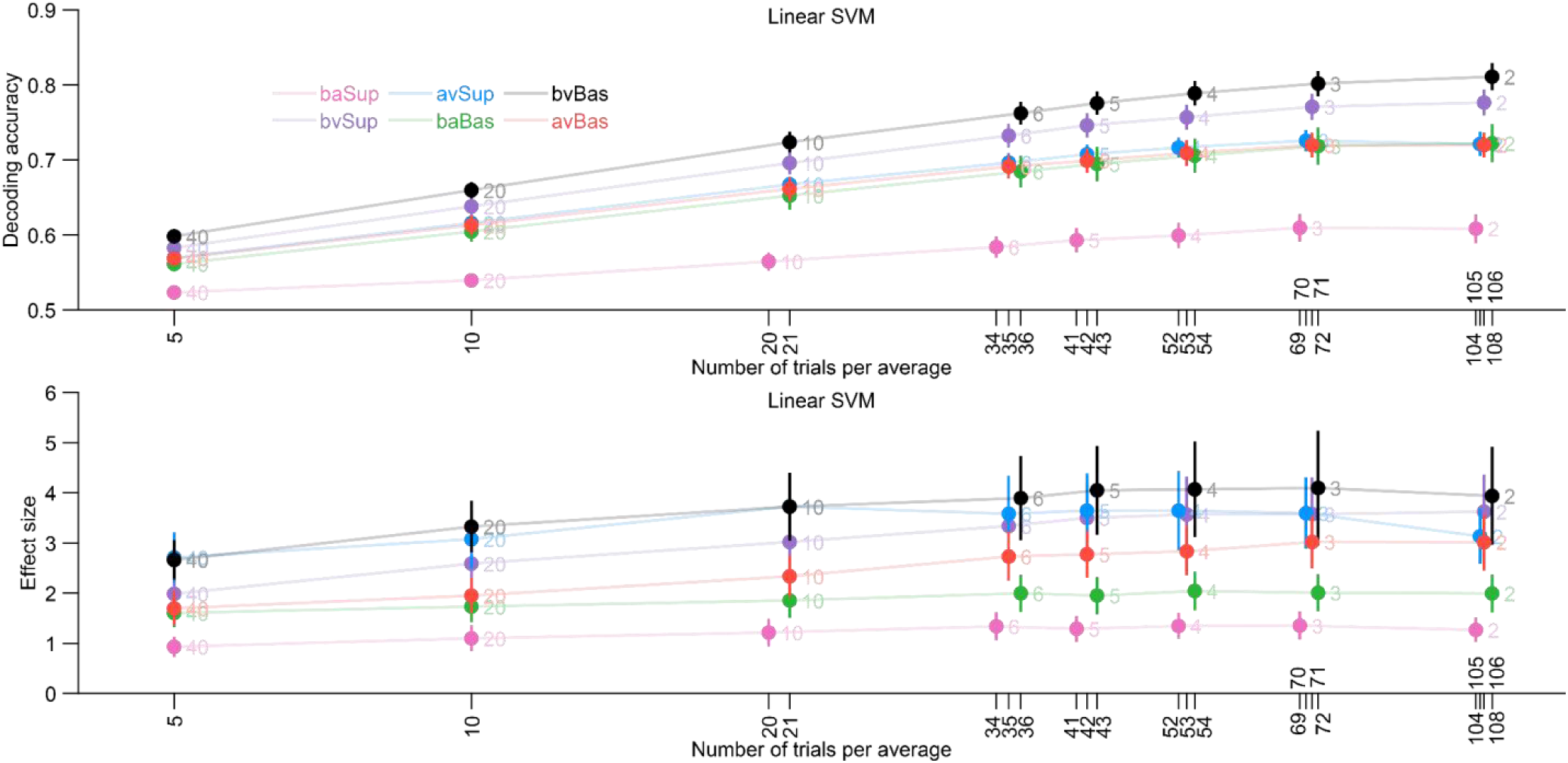
Decoding accuracy (top row) and effect size (bottom row) for different numbers of crossfolds for the dataset from Poncet et al (2025). Decoding accuracy was averaged across a time window of 100 to 500 ms after stimulus onset. The X axis labels indicate the number of trials per average, and the number next to each data point indicates the corresponding number of crossfolds. Error bars show the standard error for each value. The chance level is 0.5 for decoding accuracy and zero for effect size in all cases. baSup: Bird/Non-bird animal at the superordinate level; avSup: Non-bird animal/Vehicle at the superordinate level. bvSup: Bird/Vehicle at the superordinate level; baBas: Bird/Non-bird animal at the basic level; avBas: Non-bird animal/Vehicle at the basic level. bvBas: Bird/Vehicle at the basic level.

### S4. Effect Sizes Based on Single-Time-Point Decoding Accuracy

**Figure S10.**
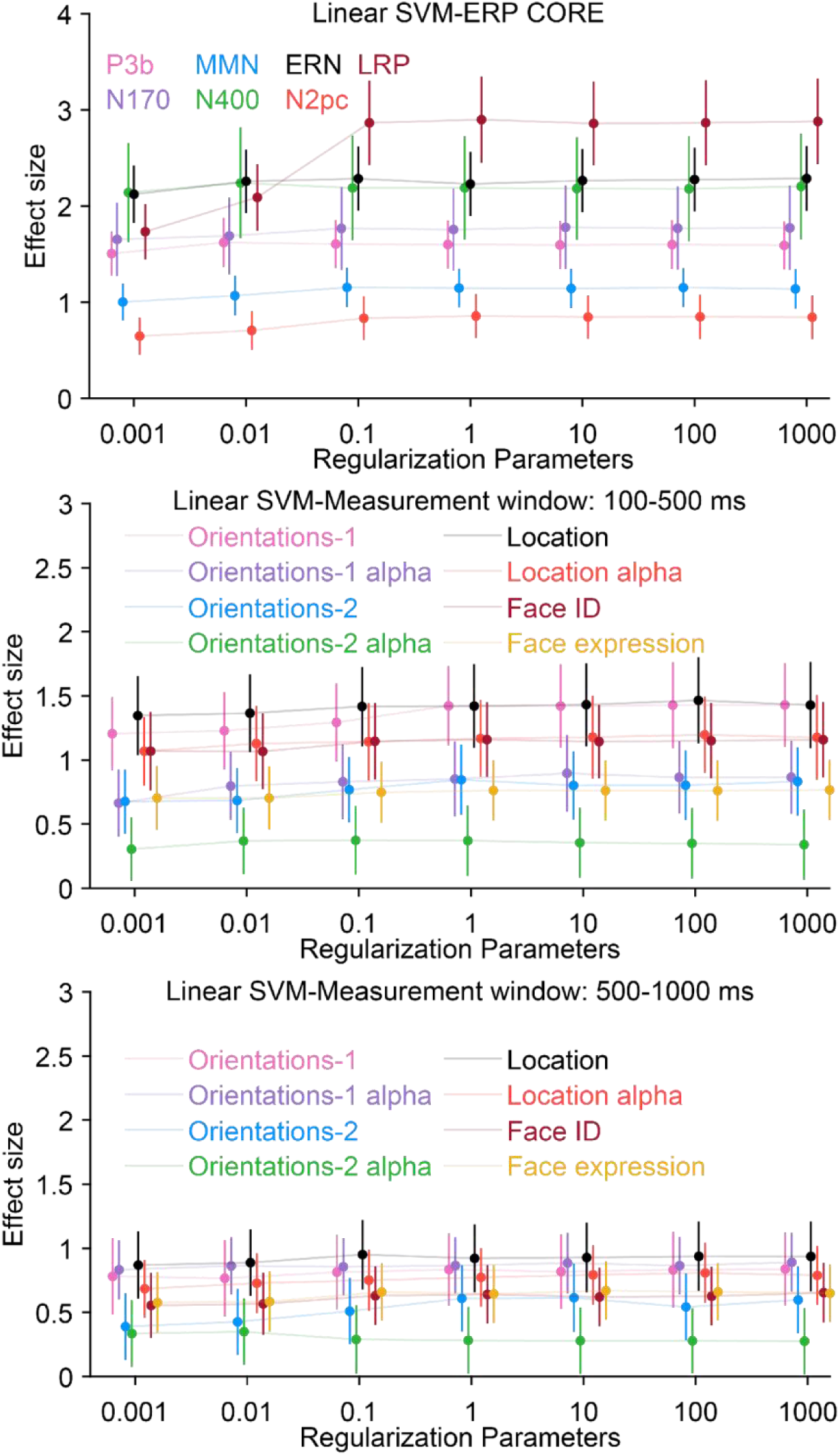
Effect size that was first computed separately at each time point and then averaged over the time range of interest for different regularization strengths, separately for each dataset. Error bars show the standard error for each value.

**Figure S11.**
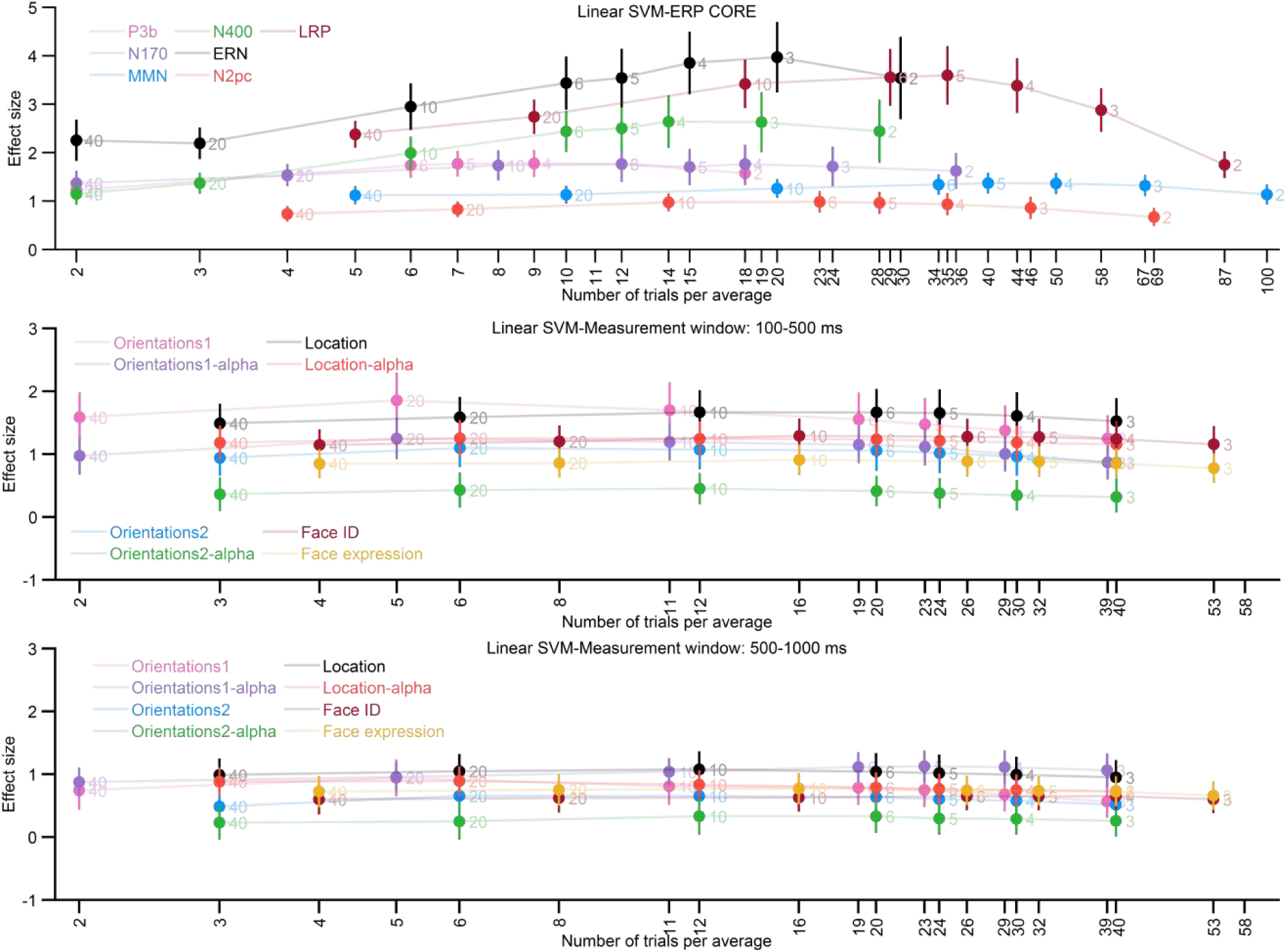
Effect size that was first computed separately at each time point and then averaged over the time range of interest for different numbers of crossfolds, separately for each dataset. Error bars show the standard error for each value.

